# Cryo-EM structures reveal interactions of S-OPA1 with membrane and changes upon nucleotide binding

**DOI:** 10.1101/528042

**Authors:** Danyang Zhang, Yan Zhang, Jun Ma, Tongxin Niu, Wenbo Chen, Xiaoyun Pang, Yujia Zhai, Fei Sun

## Abstract

Mammalian mitochondrial inner membrane fusion is mediated by OPA1(optic atrophy 1). Under physiological condition, OPA1 undergoes proteolytic processing to form a membrane-anchored long isoform (L-OPA1) and a soluble short isoform (S-OPA1). A combination of L-OPA1 and S-OPA1 are required for membrane fusion, however, the relevant mechanism is not well understood. In this study, we investigate the cryo-EM structures of S-OPA1 coated liposome at nucleotide-free and GTPγS bound states. S-OPA1 exhibits a general structure of dynamin family. It can assemble onto membrane in a helical array with a building block of dimer and thus induce membrane tubulation. A predicted amphipathic helix is discovered to mediate the tubulation activity of S-OPA1. The binding of GTPγS triggers a conformational rotation between GTPase domain and stalk region, resulting the rearrangement of helical lattice and tube expansion. This observation is opposite to the behavior of other dynamin proteins, suggesting a unique role of S-OPA1 in the fusion of mitochondrial inner membrane.

**SIGNIFICANCE STATEMENT:** Mitochondria are highly dynamic cellular organelles that constitute a remarkably dynamic network. Such dynamic network is vital to keep homeostasis of cellular metabolism and it is balanced by fission and fusion events. Having the double membrane, the fusion of mitochondria becomes more complicated in comparison with other cellular mono-membrane organelles. The inner membrane fusion is driven by OPA1 that needs to be pre-processed to long and short forms while the molecular mechanism is largely unknown. This work well characterizes the biochemical property of the short form of OPA1, and reveals how it interacts with membrane and how its conformation responds to nucleotide binding. This work gives a further insight into mitochondrial inner membrane fusion mechanism.

## INTRODUCTION

In eukaryotic cells, series of discrete membranous compartments separate different biochemical reactions, and the membrane fission and fusion mechanisms accomplish the communication between and within these compartments (McNew et al., 2013). A family of large GTPases called dynamins plays pivotal roles in the fission and fusion process (Praefcke and McMahon, 2004). Mitochondria are highly dynamic organelles and their remarkably dynamic networks are also regulated by large GTPases dependent fission and fusion of outer and inner membranes (Labbe et al., 2014; van der Bliek et al., 2013; Westermann, 2010). The GTPase OPA1 (optic atrophy 1), a member of dynamin protein family, is known related to mitochondrial inner membrane fusion (Anand et al., 2014; Frezza et al., 2006; MacVicar and Langer, 2016).

OPA1 composes an N-terminal mitochondrial targeting sequence (MTS), a following transmembrane domain (TM), a coiled coil domain, a highly conserved GTPase domain, a middle domain and a C-terminal GTPase effector domain (GED), and it has eight different spliced variations at the region between the TM and coiled coil domain (Belenguer and Pellegrini, 2013; Olichon et al., 2007) (see also **Figure 1A**). The GTPase domain, middle domain and GED are classical dynamin regions. After imported into mitochondria, the MTS is proteolytically processed to form a membrane anchored long form L-OPA1, and the L-OPA1 can be further cleaved into a short form S-OPA1 through the S1 or S2 site between the TM and coiled coil domain (Naotada Ishihara et al., 2006; Song et al., 2007). Both the long and short form of OPA1 participate in mitochondria inner membrane fusion, however it remains open about the specific role of S-OPA1 during the fusion procedure (Anand et al., 2014; Ban et al., 2017; Del Dotto et al., 2017).

**Figure 1.**
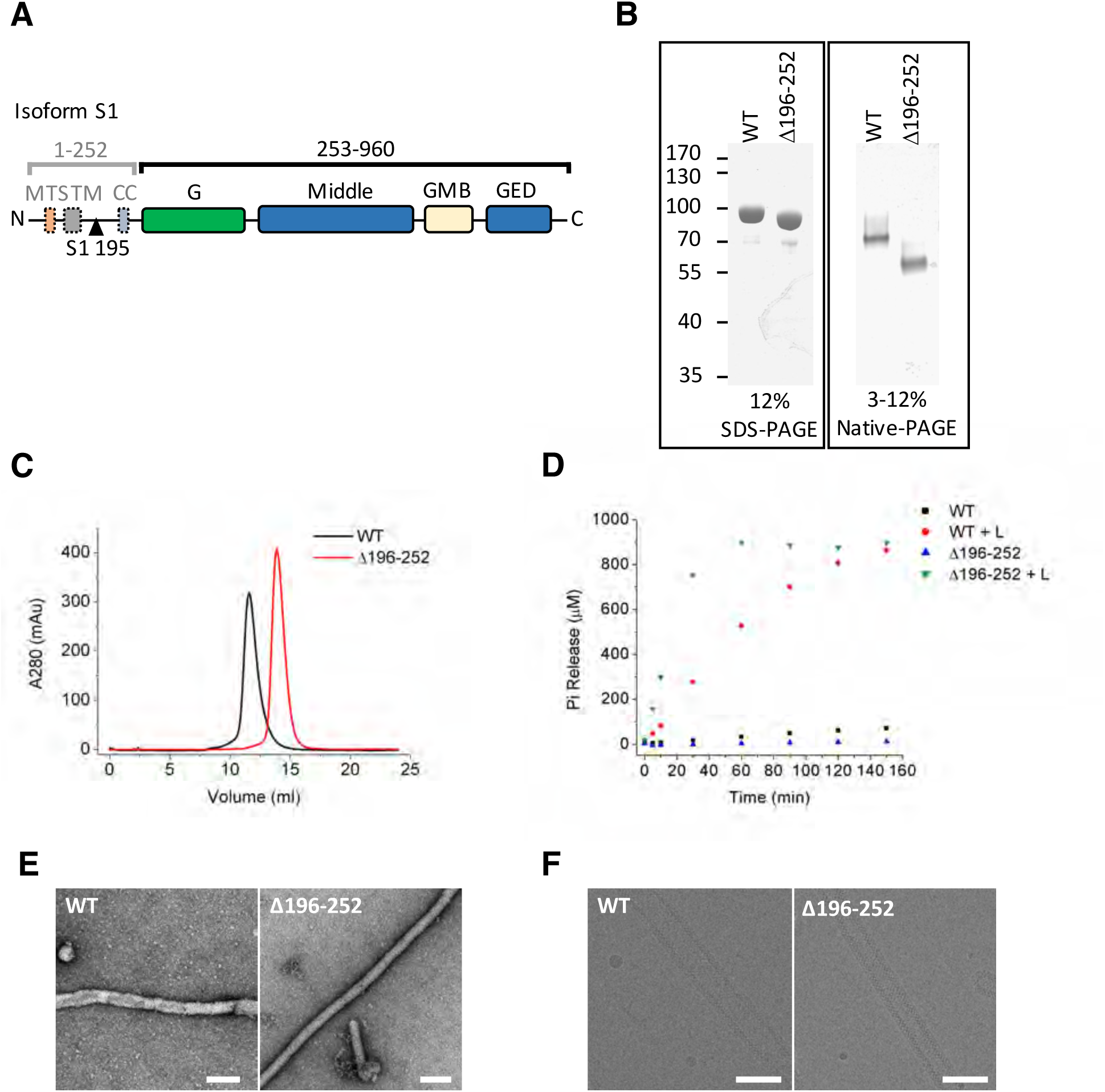
Purification and characterization of S-OPA1. (**A**) Domain organization of OPA1. MTS, mitochondrial targeting sequence; TM, transmembrane region; G, G domain; Middle, middle domain; GMB, global membrane binding domain; GED, GTPase effector domain. The proteolytic cleavage site S1 in isoform 1 at the 195th amino acid is indicated by black triangle. (**B**) SDS-PAGE and native PAGE of wild type S-OPA1 and its truncation form (Δ196-252). (**C**) Size exclusion chromatography of S-OPA1 using Superdex 200 10/300 GL column (GE Healthcare). The elution volume of the column was pre-calibrated using standard protein molecular weight markers. (**D**) Base and liposome-binding induced GTPase activity of S-OPA1 and Δ196-252. The total free phosphate was measured at each time point and data presented come from 3 independent experiments. (**E**) Negative stain electron micrographs of S-OPA1 coated tubes, scale bar: 200 nm. (**F**) Cryo electron micrographs of S-OPA1 coated tubes. Scale bar: 100 nm.

Efforts to recapitulate the fusion mechanism in vitro using FRET suggested that L-OPA1 alone on either side of membrane can promote fusion with a proper concentration of cardiolipin in the opposite membrane (Ban et al., 2017). The S-OPA1, on the other hand, bridges the opposite membrane probably via interactions with both L-OPA1 and cardiolipin and assists the L-OPA1-dependent fusion with the needs of GTP hydrolysis (Ban et al., 2017). Studies on Mgm1, the yeast homolog of OPA1, have drawn a similar conclusion in which its long form L-Mgm1 acts as a fusion-prone protein with its GTPase activity inhibited and its short form S-Mgm1 drives fusion procedure through GTP hydrolysis (DeVay et al., 2009; Zick et al., 2009). Another study of S-OPA1 confirms its tubulation activity with cardiolipin containing liposomes by negative staining electron microscopy (Ban et al., 2010). These studies speculate a GTPase dependent auxiliary function of S-OPA1 while membrane fusion, however, there are also other reports supporting the fission favorable function of S-OPA1 (Anand et al., 2014).

To further understand the role of S-OPA1 in mitochondrial inner membrane fusion, here we substantially studied the biochemical property of S-OPA1 and utilized cryo-electron microscopy to solve the structures of S-OPA1 coated liposome in a nucleotide-free state and a GTPγS binding state. Our studies imply a distinctive role of S-OPA1 in mitochondrial inner membrane fusion.

## RESULTS

### S-OPA1 can induce tubulation of cardiolipin containing liposome

We expressed the short isoform of splice form 1 human OPA1 (S-OPA1, see **Figure 1A**) in bacteria and purified into homogeneity (**Figure 1B**). Gel-filtration and chemical cross-linking experiments indicate a dimerization form of S-OPA1 (**Figure 1C; see also Figures S1A and S1B**). The GTP hydrolysis activity of S-OPA1 is weak but significantly enhanced ∼70 fold (Kcat) with the existence of liposome (**Figure 1D and Table S1**). The liposome was prepared with a phospholipid composition of 45% DOPC (1,2-dioleoyl-sn-glycero-3-phosphocholine), 22% DOPE (1,2-dioleoyl-sn-glycero-3-phosphoethanolamine), 8% PI (phosphatidylinositol) and 25% CL (cardiolipin), which approximates the composition of mitochondrial inner membrane (Ban et al., 2010). By examining the mixture of S-OPA1 with liposome using both negative electron microscopy (nsEM) and cryo-electron microscopy (cryo-EM), we found significant tubulation of liposome induced by S-OPA1 (**Figures 1E and 1F**). However, we found the tubes are varied with the diameters and diffraction patterns, suggesting the membrane coated S-OPA1 did not form a unique helical lattice.

To optimize the homogeneity of the sample, we cloned and expressed a truncation form of S-OPA1 (S-OPA1-Δ196-252, see **Figure 1A and 1B**) by deleting its N-terminal residues from 196 to 252 because this region was reported responsible for the dimerization of S-OPA1 in solution (Akepati et al., 2008). Compared to the wild type, the truncation form behaves as a monomer in gel-filtration (**Figure 1C and S1B**) and its basal GTP hydrolysis activity is lower but can be stimulated about two orders with the existence of liposome (**Figure 1D and Table S1**). In addition, the truncation form can induce homogeneous tubes with smaller diameter (**Figures 1E and 1F**). If not specially mentioned, all the subsequent cryo-EM structural studies were performed using this truncation form S-OPA1-Δ196-252. However, all the subsequent biochemical and biophysical assays were performed using the wild type full length S-OPA1.

### Helical structure of S-OPA1 coated liposomal tube in a nucleotide-free state

To characterize the structure of nucleotide-free S-OPA1 coated liposomal tube, we collected a cryo-EM dataset of S-OPA1 coated tubes and classified the boxed tubes via the diameters and diffraction patterns (**Figures S1C and S1D**). A selected class of tubes with the average diameter of 53 nm were segmented and further reconstructed by the iterative helical real-space reconstruction (IHRSR) algorithm (Egelman, 2000, 2007) (**Figure S1E and S1F**). This approach finally generated a six-start left-handed helical map at a resolution of ∼ 14 Å. The map has an inner diameter of 23 nm, an outer diameter of 53 nm, 17.3 units per turn and a pitch of 465.6 Å (**Figures 2A, 2B and Movie S1**).

**Figure 2.**
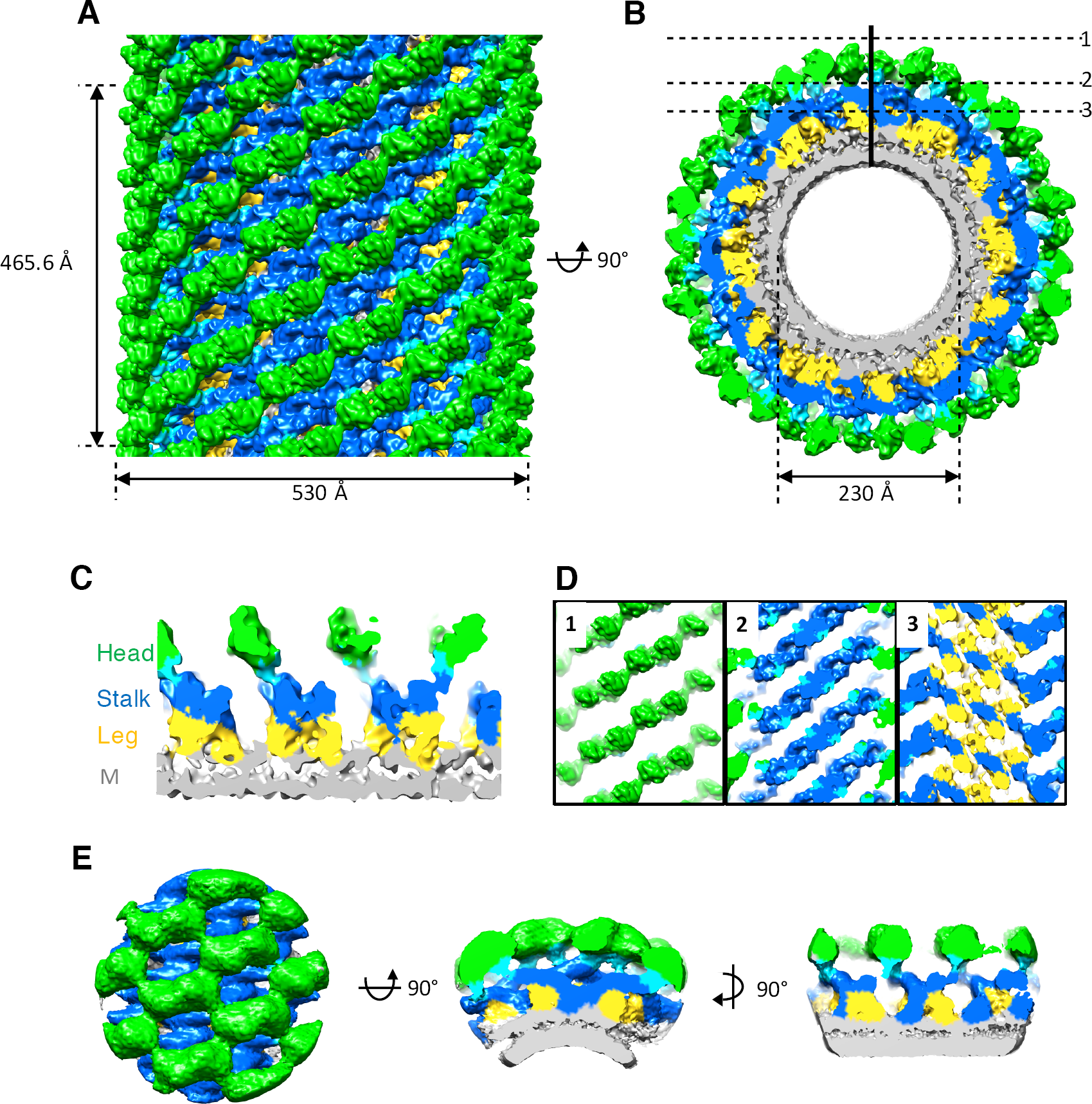
3D reconstruction of nucleotide-free S-OPA1 coated tubes. (**A**) Side view of cryo-EM map of S-OPA1 coated tube. Other than membrane, the map is subdivided and colored radially into three layers denoting “leg” (yellow), “stalk” (blue), and “head” (green and cyan). The outer diameter and pitch are labeled. (**B**) Radical cross-section of the tube. The inner diameter is labeled. Dashed black lines denote the planar sections that are rotated by 90° and shown in (D). (**C**) Cross-section of the tube along the solid vertical black line in (B). The leg, stalk, head, and membrane bilayer density are labeled and colored as in (A). (**D**) Corresponding cross sections of the tube along the dashed black lines in (B). The density color scheme is same with (A). **(E)** Sub-volume averaging of S-OPA1 coated tube at nucleotide-free state. The map was viewed at the side (left panel), end-on (middle panel) and section (right panel) as (A), (B) and (C), correspondingly.

Similar to the dynamin 1 (Dyn1) coated tube (Chappie et al., 2011; Sundborger et al., 2014), the S-OPA1 coated tube can be generally divided into three regions along its radial direction, an inner density fused with the outer leaflet of lipid bilayer, a middle density packing compactly, and an outer globular density (**Figures 2C and 2D**). Thus we name the inner, middle, and outer density as leg, stalk and head respectively in consistence with the nomenclature of Dyn1 coated tube (Chappie et al., 2011).

To be noted, since we could not determine the handedness of S-OPA1 coated tube at the current resolution from helical reconstruction, we further performed cryo-electron tomography (cryo-ET) combined with sub-volume averaging (SVA) to determine the structure of S-OPA1 coated tube (The handedness of this procedure has been pre-calibrated). The final averaged cryo-ET map shows a consistent architecture **(Figure 2E and S2A**) and corrects the handedness of the helical reconstruction (**Figure 2**). We also determined the structure of full length S-OPA1 coated tube by using the same tomographic procedure (**Figure S2B**), showing the same architecture with the truncation form. Thus, our observation of the helical lattice of S-OPA1 bound to membrane is not an artifact from the truncation of the coiled coil domain.

### Domain organization of S-OPA1 and membrane binding sites

Dynamin proteins have similar domain architecture (**Figure 1A and 3A**), and structure predictions by Phyre2 (Kelley et al., 2015) and I-TASSER (Roy et al., 2010; Yang et al., 2015; Zhang, 2008) showed that S-OPA1 has a classic Dyn1-like general structure (**Figure S1G**). Since S-OPA1 shares the highest degree of sequence conservation (19.8% identical and 35.0% similar) with Dyn1, we used the crystal structure of human Dyn1 (PDB ID 3SNH) (Faelber et al., 2011) to dock into the cryo-EM map. Considering potential relevant conformational changes among different domains, we separated the crystal structure of Dyn1 into three parts, the G/BSE region, the middle/GED stalk, and the PH domain (**Figure 3A**).

**Figure 3.**
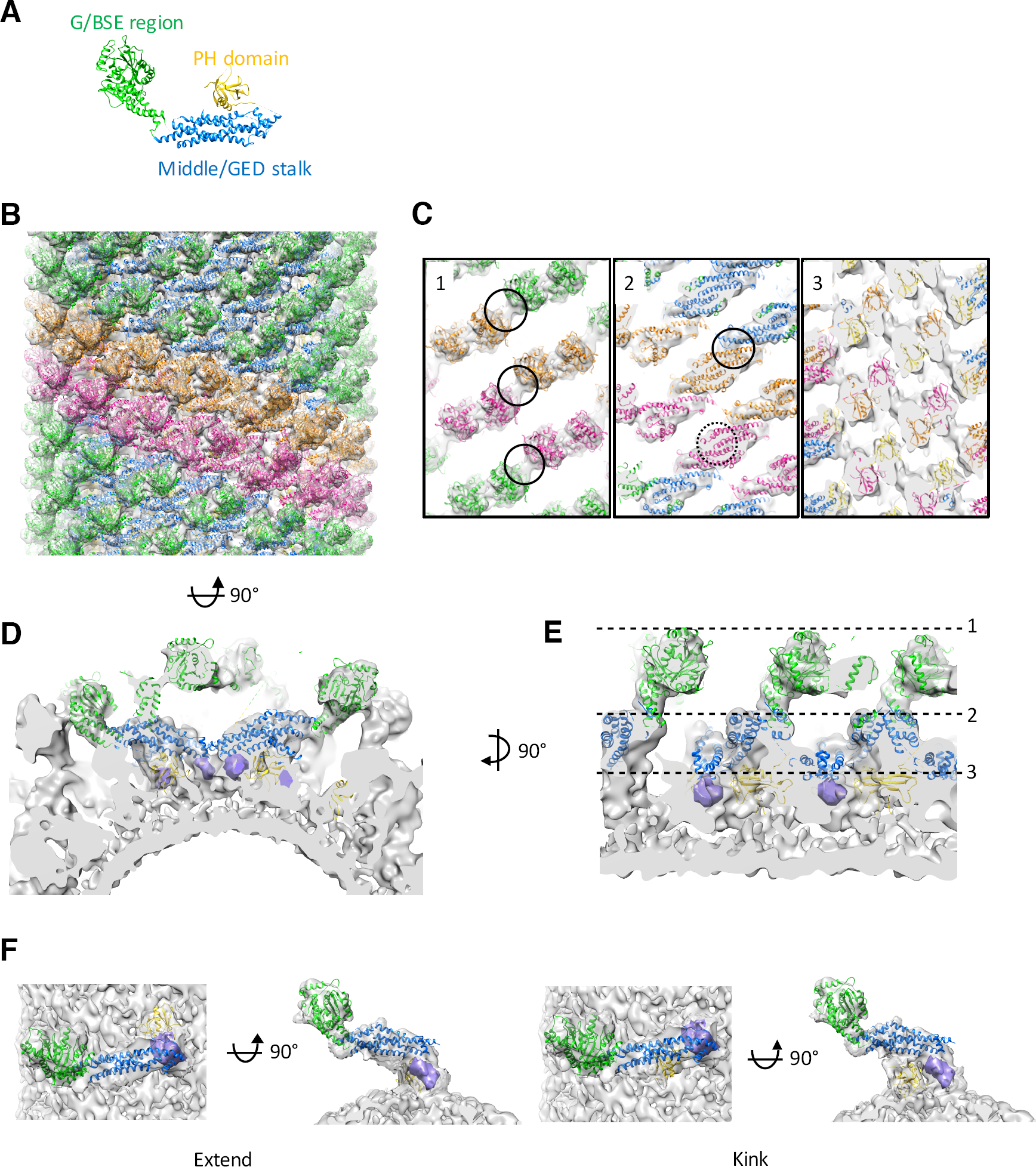
Docking crystallized fragments of dynamin1 into cryo-EM map of nucleotide-free S-OPA1 coated tube. (**A**) Crystal structure of nucleotide-free human Dyn1 (Faelber et al., 2011) (PDB ID 3SNH). The structure is separated into G/BSE region, middle/GED stalk region and PH domain, respectively. (**B**) Docking structural fragments of nucleotide-free human Dyn1 into cryo-EM map (transparent gray) of nucleotide-free S-OPA1 coated tube. G/BSE domains are colored green, and middle/GED stalk are colored blue. Building blocks in two different helical starts are colored orange and pink, respectively. (**C**) Slicing cross sections of the tube showing the fitness between structural model and the map. The positions of cross sections are same with **Figure 2D** and (E). Black circles in panel 1 indicate potential G dimer interface. Solid line circle in panel 2 highlights bundle-to-bundle stalk interface and dashed line circle for the tip-to-tip stalk interface. (**D**) Zoomed–in view of radical cross-section showing the fitness between structural model and the map. The PH domain of human dynamin 1 is fitted into the leg density and the density belonging to the extension of the stalk is shown in a higher threshold and colored in purple. (**E**) Vertical cross section of the map that rotates 90° with respect to (D). Dashed black lines denote the positions of the cross sections in (C). (**F**) Two possible monomer conformations of S-OPA1, the extended form and the kinked one.

The G/BSE region can be well fitted into the head layer and the linker density that connects the head and stalk layers (**Figures 3B, 3D and 3E**). The good fitness suggests S-OPA1 has a similar structural component of G domain (GTPase domain) and BSE three-helix bundle with Dyn1. Since the G domains of dynamin proteins need to dimerize to activate the GTPase activity (Chappie et al., 2010; Raphael Gasper et al., 2009), we also investigated whether the G domains of S-OPA1 are dimerized in the present cryo-EM density. However, the attempts to dock the crystal structure of dimerized G domains GG_GDP.AlF4_-(PDB ID 2X2E) (Chappie et al., 2010) failed (**Figure S3A**). And the dimerization interfaces of the proximal G domains in the present packing are separated with about 30 angstroms (**Figure 3C, 1**), suggesting a further conformational change is needed in the subsequent nucleotide binding to enable the activity of GTP hydrolysis.

The middle/GED stalk structure can be well docked into the stick-like density of the middle/stalk layer of the map (**Figures 3B, 3C and 3D**). However, the stalk layer of S-OPA1 may have a slightly different conformation compared to the middle/GED stalk structure of Dyn1 as its distal tip stretches into the inner layer and directly interacts with the membrane (**Figures 3D, 3E and S3B**). The whole stalk region of the S-OPA1 array shows a compact packing with two interaction interfaces, the bundle-to-bundle interface and the tip-to-tip one (**Figure 3C, 2**). Such compact packing suggests a pivotal role of stalk interactions of S-OPA1 in maintaining the structural stability of the protein-lipid complex.

The leg density of the S-OPA1 tube is more complicated than the Dyn1 tube (Chappie et al., 2011) because it not only contains a globular density corresponding to the leg of Dyn1 but also contains another density corresponding to the extension of S-OPA1 stalk (**Figures 3C, 3D, 3E and S3B**). Sequence analysis does not suggest a membrane binding domain of S-OPA1 explicitly, however, we were surprised to find the Dyn1 PH domain could fit into the globular density very well (**Figures 3C, 3D and 3E**), suggesting S-OPA1 would have a globular membrane binding domain (GMB) with a comparable size of PH domain and this GMB domain interacts with the mitochondrial inner membrane. The corresponding sequence of such domain would be located between the middle and GED domain (**Figures 1A and S5A**).

The current resolution of the map could not determine the linkage between the stalk and the GMB domain. However, according to the packing of S-OPA1 on the tube, there are only two possible conformations, a kinked conformation with the GMB domain right underneath its own stalk region or an extended conformation with the GMB domain stretching into the nearby interstice (**Figure 3F**).

Overall, based on the present cryo-EM density of S-OPA1 coated tube, we observed two membrane binding sites of S-OPA1, one is located at its GMB domain and another at the extension of its stalk domain (**Figure S3B**).

### S-OPA1 dimer is the building block of its helical array on liposomal tube

The helical reconstruction imaging processing procedure indicated S-OPA1 coated tube has a 6-fold symmetry and contains six helical starts. By fitting crystal structures of Dyn1 domains, we found the asymmetric unit of S-OPA1 packing array contains two copies of S-OPA1 molecules that form a dimer (blue and gray in **Figure S3C**). There are two possible forms of S-OPA1 dimer according to the packing array, one is called here the short dimer (green in **Figure S3C**) and the other defined as the long dimer (pink in **Figure S3C**). The short dimer utilizes the tip-to-tip interactions of their stalk domains to form the dimerization interface while the long dimer is more extended and its dimerization interface is mediated by their stalks bundle-to-bundle interactions. Our subsequent structural analyzes by comparing structures of S-OPA1 coated tubes in different nucleotide binding states suggest the short dimer is more likely the building block of S-OPA1 assembly on liposome (see context below and also **Figure S8A**). Thus, in the following structural analysis and comparison, we will only focus on the short dimer.

We then investigated how the S-OPA1 dimers assemble on the liposomal tube. By following the helical symmetry, the S-OPA1 short dimers assemble into a single helical turn and then form one start of the packing array (**Figure 3B**). However, for each start, the building blocks are not close enough to form interactions. The packing array needs to be stabilized by the interactions among different starts and we found such interactions are mediated by bundle-to-bundle interfaces of stalks (**Figures 3B and 3C**). These stalk interactions are the main factors to stabilize the S-OPA1 helical structure. In addition to the stalk interactions, we noticed the G domains of S-OPA1 in adjacent helical starts are facing each other with their dimerization interfaces opposed, which leaves a potential for the subsequent change of S-OPA1 array (**Figure 3C, 1**).

### S-OPA1 membrane tubulation activity is independent with its GTPase activity

Our observation that S-OPA1 can induce tubulation of liposome without the addition of nucleotide has suggested the tubulation activity of S-OPA1 is independent with its GTP hydrolysis activity. To further validate this assumption, we performed mutagenesis. The high sequence conservation of G domains among dynamin proteins enabled us to identify the locations (a.a. 297-302 and 319-327) of the key catalytic residues of S-OPA1 (**Figure S4A**) at its P loop and switch I (Schmid and Frolov, 2011). All the mutants Q297E, S298A, G300E, T302N and T323A kept their wild-type abilities to bind to liposome but lost their liposome induced GTPase activities (**Figures 4A, 4B and Table S1**). However, interestingly, their tubulation activities did not change significantly by examining their liposomal tubes using nsEM (**Figure 4C**). These results confirm S-OPA1 can deform membrane without the need of GTP hydrolysis and further suggest the GTP hydrolysis of S-OPA1 occurs most likely after the liposomal tubulation.

**Figure 4.**
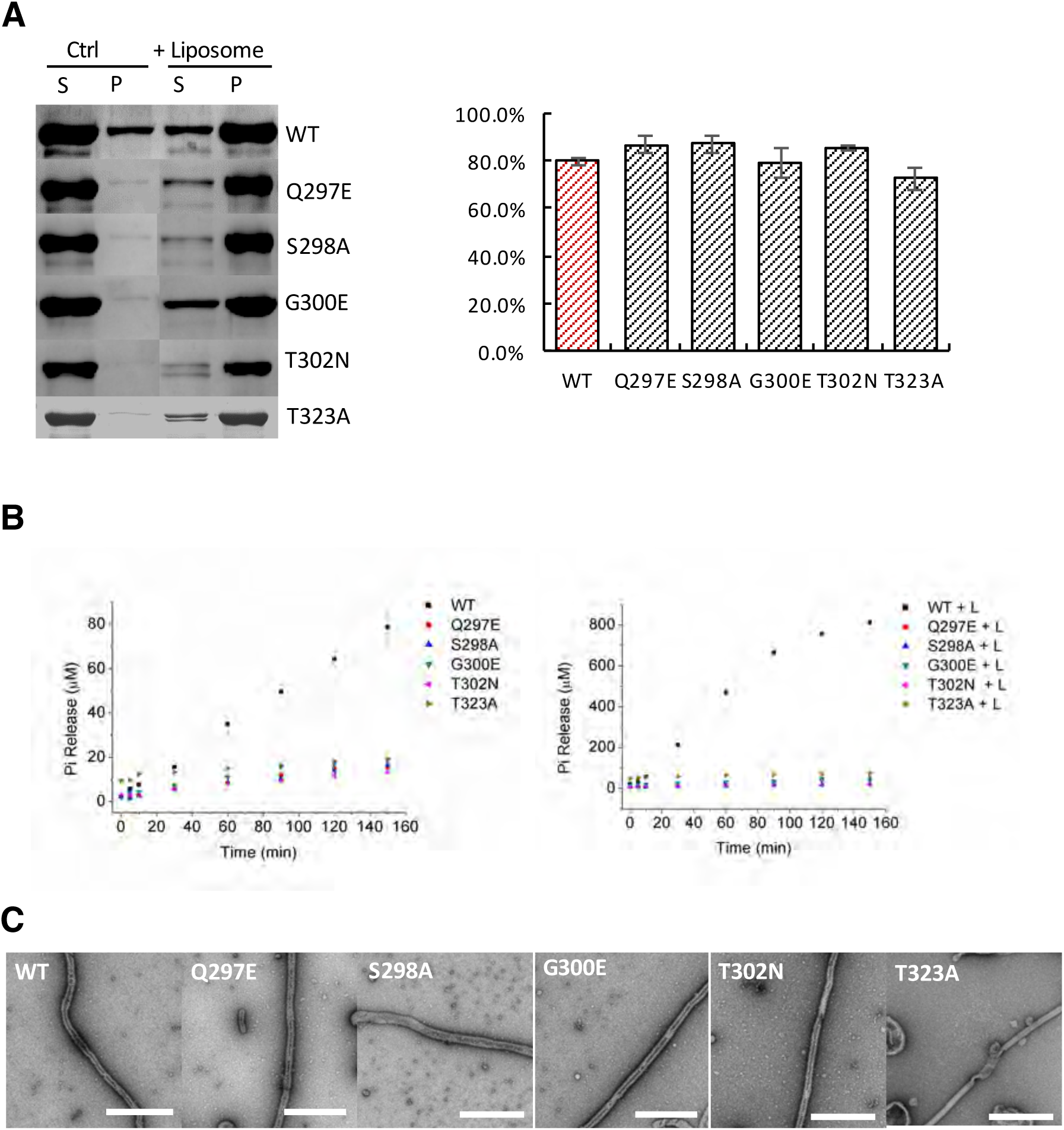
Tubulation activity of S-OPA1 is independent with its GTPase activity. (**A**) Sedimentation of wild type S-OPA1 and its G domain mutants with or without cardiolipin containing liposomes (n=3). S, supernatant; P, pellet; *, P<0.01; **, P<0.001. (**B**) Basel (left panel) and liposome binding induced (right panel) GTPase activity of wild-type S-OPA1 and its G domain mutants. L, liposome. The total free phosphate was measured at each time point and data presented result from 3 independent experiments. (**C**) Tubulation activity of wild-type S-OPA1 and G domain mutants examined by negative stain electron microscopy. Scale bar: 500 nm.

### S-OPA1 has a potential amphipathic helix involved in membrane tubulation

Next, we sought to identify the factors to affect S-OPA1 membrane tubulation activity. The above structural analysis suggests the GMB domain of S-OPA1 would be presumably responsible for S-OPA1 membrane binding and deformation. Sequence analysis suggests such GMB domain would correspond to the residues from 734 to 850 in S-OPA1 (**Figure S5A**). Subsequent secondary structure prediction and helical wheel analysis (Alan Bleasby, pepwheel, http://www.bioinformatics.nl/cgi-bin/emboss/pepwheel) identified a possible amphipathic helix between E794 and K800 (**Figure S5B**). Previous studies have discovered the important role of an amphipathic helix in membrane remodeling (Shen et al., 2012). We thus performed mutagenesis to investigate this susceptible amphipathic helix in S-OPA1 GMB domain for its role in membrane tubulation.

We constructed five mutants of S-OPA1 by mutating the region 794-800 to all alanine residues (794-800A), or by point mutations (E794AE795A, K797AK800A, L795EM798EL799E and L795AM798AL799A). Compared to wild type, the five mutants exhibited moderate suppression on membrane binding suggesting this region partially involves in membrane binding via hydrophobic interaction (**Figure 5A**). However, we observed more significant effects of these mutants upon their liposome induced GTPase activity (**Figure 5B and Table S1**) and membrane tubulation activity (**Figure 5C**). The mutants 794-800A, L795EM798EL799E and L795AM798AL799A completely lost their liposome induced GTP hydrolysis activities and membrane tubulation activities. While, these activities of mutants E794AE795A and K797AK800A were also greatly reduced. As a result, the predicted amphipathic helix of GMB domain would be one of the major factors involved in S-OPA1 induced membrane tubulation procedure, which most likely inserts hydrophobic residues into the membrane (Drin and Antonny, 2010). The reduced or abolished membrane tubulation activities therefore affect the subsequent membrane induced GTP hydrolysis activity.

**Figure 5.**
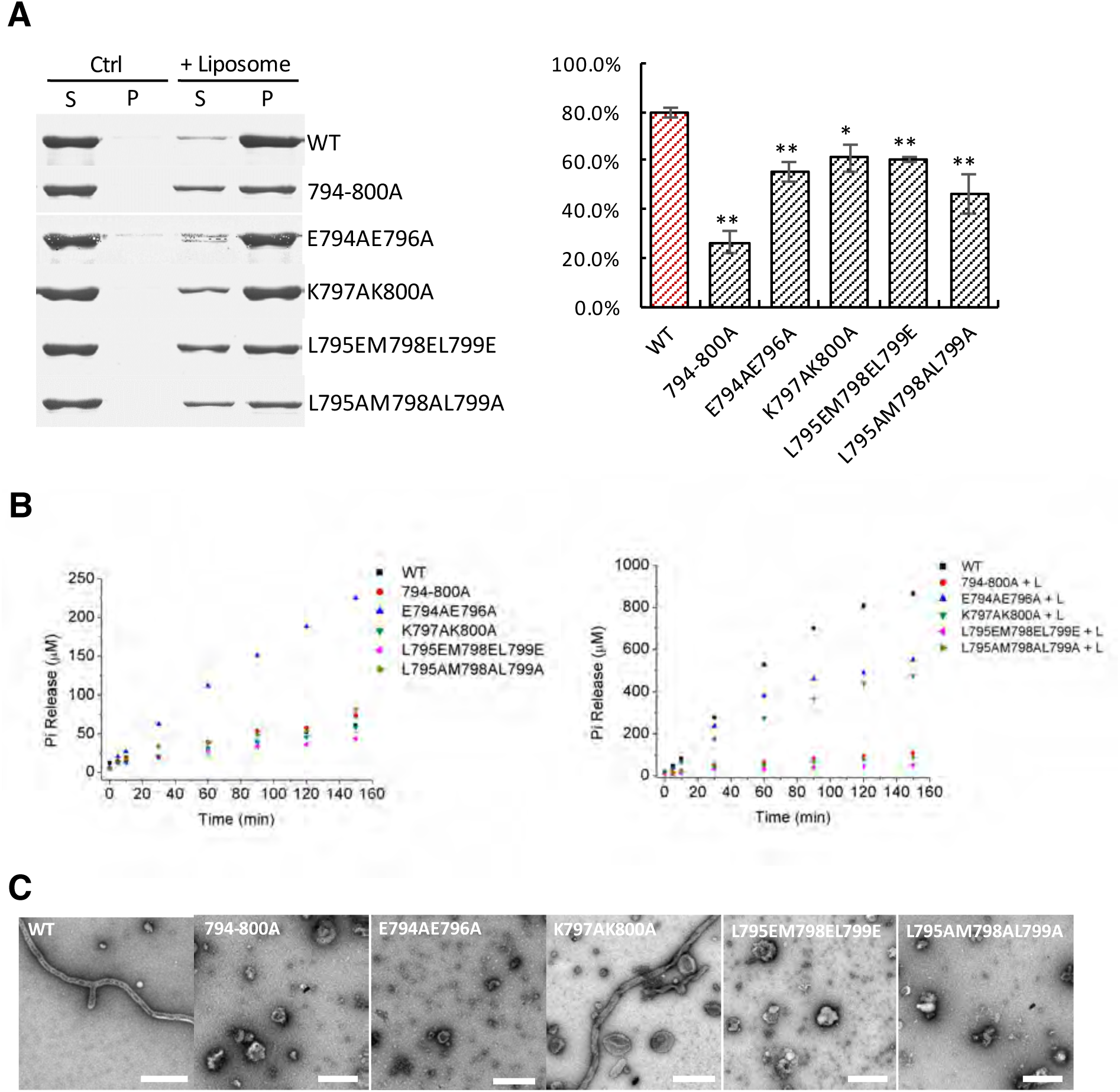
A predicted amphipathic helix in S-OPA1 is indispensable for its tubulation activity. (**A**) Sedimentation of wild type S-OPA1 and its mutants with or without cardiolipin containing liposomes (n=3). S, supernatant; P, pellet; *, P<0.01; **, P<0.001. (**B**) Basel and liposome binding induced GTPase activity of wild-type S-OPA1 and its amphipathic helix mutants. L, liposome. The total free phosphate was measured at each time point and data presented come from 3 independent experiments. (**C**) Tubulation activity of wild-type S-OPA1 and its mutants examined by negative stain electron microscopy. Scale bar: 500 nm.

In addition, the above cryo-EM map based structural analyzes have identified two membrane binding sites of S-OPA1. The mutagenesis studies here indicated a less important role of the predicted amphipathic helix of the GMB domain in membrane binding. Therefore, another identified stalk tip region of S-OPA1 would play a more important role for S-OPA1 bound to membrane.

### Nucleotide binding induces a reduced curvature of S-OPA1 coated tube

Since the GTPase activity of S-OPA1 is indispensable for its fusion in promoting mitochondrial inner membrane fusion (Ban et al., 2010), we sought to study how nucleotide binding could take effect on the S-OPA1 coated liposomal tube. Significant conformational changes of dynamin proteins after nucleotide binding have been observed for Dyn1 and Dnm1 (Chappie et al., 2011; Frohlich et al., 2013; Mears et al., 2011).

We incubated excess GTP or its non-hydrolyzed or slowly hydrolyzed analogs (GMPPCP, GMPPNP and GTPγS) with the wild type and truncated S-OPA1 coated liposomal tubes for 30 mins and examined the tubes by cryo-electron microscopy (**Figure S6A**). We surprisingly found that the diameters of the tubes were all increased from the original ∼ 53 nm to 70∼80 nm (**Figure S6B**), which is in contrast to the phenomena observed for Dyn1 and Dnm1, where a constricted diameter was observed after nucleotide binding (Chappie et al., 2011; Frohlich et al., 2013; Mears et al., 2011). Besides the expansion of the tube diameter and the reduction of the tube curvature, compared to the nucleotide-free state, the binding of nucleotide also decreases the homogeneity of the S-OPA1 coated tubes with a wide range of tube diameters (**Figure S6B**), suggesting an increased variability of S-OPA1 in the nucleotide bound state.

We observed the instability of the S-OPA1 coated liposomal tubes after incubating with GTP, which might be due to the subsequent GTP hydrolysis. Thus, we selected GTPγS bound state (truncated S-OPA1 coated) for the subsequent structural studies because this sample is stable enough and relative homogenous in comparison with other states. Helical reconstruction technique failed due to the variable diameters. We therefore utilized cryo-ET and SVA to analyze the assembly of GTPγS bound S-OPA1 on the liposomal tube (**Figures S6C, S6D, S6E and S6F**).

### A loosened helical lattice of S-OPA1 coated tube after nucleotide binding

Compared to the nucleotide-free state, the 23 Å cryo-EM map (see **Figure S6F**) of S-OPA1 coated tube with GTPγS bound also shows four layers of densities, the outer head region, the middle stalk region, the inner leg region, and the innermost membrane region (**Figures 6A and 6B**). However, after GTPγS binding, the interlaced stalks of S-OPA1 rotate ∼20° clockwise and the distance between neighboring stalk rungs expand ∼55 Å leaving significant gaps, which results in a reduced compactness of S-OPA1 assemble and a loosened helical lattice (**Figure 6A**).

**Figure 6.**
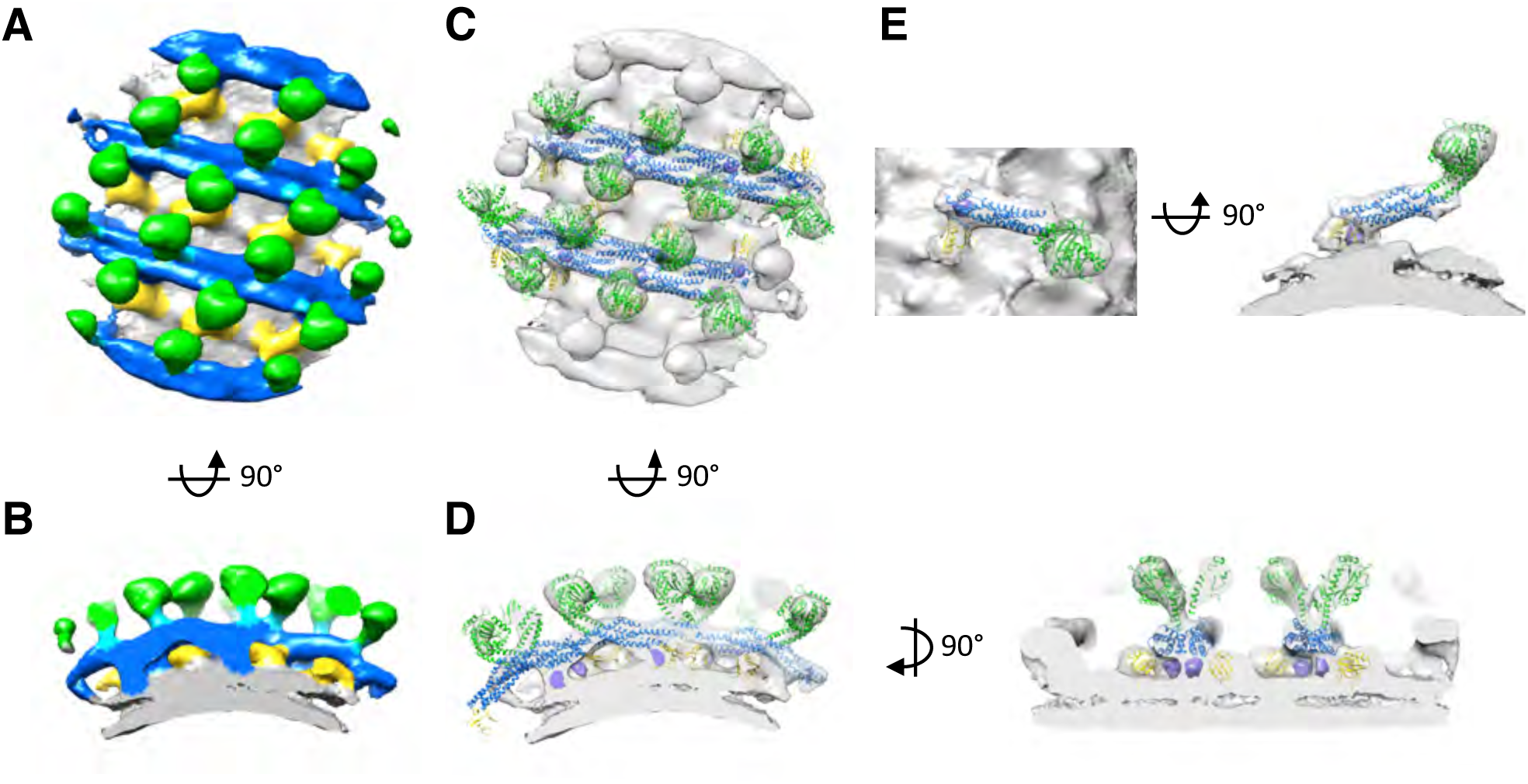
Sub-tomogram averaging of S-OPA1 coated tube in GTPγS bound state. (**A**) Side view of S-OPA1 coated tubes after adding GTPγS. The map is subdivided into three layers and colored with the same scheme in **Figure 1**. (**B**) Cross-section view of the map that is rotated 90° from (A). (**C**) Docking structure fragments of dynamin 1 into the map in (A). (**D**) Cross section views of the map that is rotated 90° from (C). The densities belonging to the extension of the stalk are shown in a higher threshold and colored in purple. (**E**) Possible monomer conformation of S-OPA1 in an extended form.

We then docked the aforesaid crystal structures of Dyn1 domains into the cryo-EM density (**Figure 6C**). The structure of G/BSE domain can be well fitted into the head region (green in **Figures 6C and 6D**) and the middle/GED domain well fitted into the middle stalk region (blue in **Figure 6C and 6D**). The potential dimerization interfaces of S-OPA1 G domains still exist between neighboring rungs (**Figure 6C**) while the distance between the G domains becomes closer (compared to nucleotide-free state) but still not close enough to form dimer **(Figure S7A)**, indicating further conformation adjustment needed for the subsequent GTP hydrolysis.

For the inner leg region, there are two pieces of densities. One (purple in **Figure 6D**) is underneath the stalk region and would account for the extension of the stalk tip of S-OPA1 for membrane interaction. Another globular density (yellow in **Figure 6A**) lies underneath the gap between the parallel stalks, which can be well fitted with a dimer of PH domain of Dyn1 (**Figures 6C and 6D**) and thus presumably account for the dimerized GMB domains. The above structural analyzes of S-OPA1 in nucleotide-free state suggest two possible conformations of S-OPA1, the kinked and the extended ones with two possible locations of its GMB domain (**Figure 3F**). However, the cryo-EM density here suggests after nucleotide binding the GMB domain of S-OPA1 deflects to the stalk flank and stays away from the stalk. Thus, an extended configuration of S-OPA1 monomer is more possible (**Figure 6E**).

By investigating the packing array of S-OPA1, we also found that the asymmetric unit of S-OPA1 coated tube after GTPγS binding contains two copies of S-OPA1 molecules, which can form either a short dimer or a long dimer (**Figure S7B**). In consistency with the above nucleotide-free one, the short dimer with GTPγS bound utilizes the same interaction interfaces (**Figure 7B**). While the long dimer in the GTPγS bound map has a completely different interface with the one in the nucleotide-free state (**Figure S8A**). Thus, by combining two cryo-EM maps of S-OPA1 coated tube in both nucleotide-free and bound state, we concluded the short form of S-OPA1 dimer is the building block when it binds to membrane and starts to assembly. We further compared this dimer of S-OPA1 with other dynamins dimers (Dyn1, MxA and Drp1) that are involved in membrane fission and found their significant differences (**Figure S8B**). The S-OPA1 utilizes the stalks tip-to-tip interaction for dimerization while other dynamins utilize the bundle-to-bundle interaction of stalks for dimerization.

**Figure 7.**
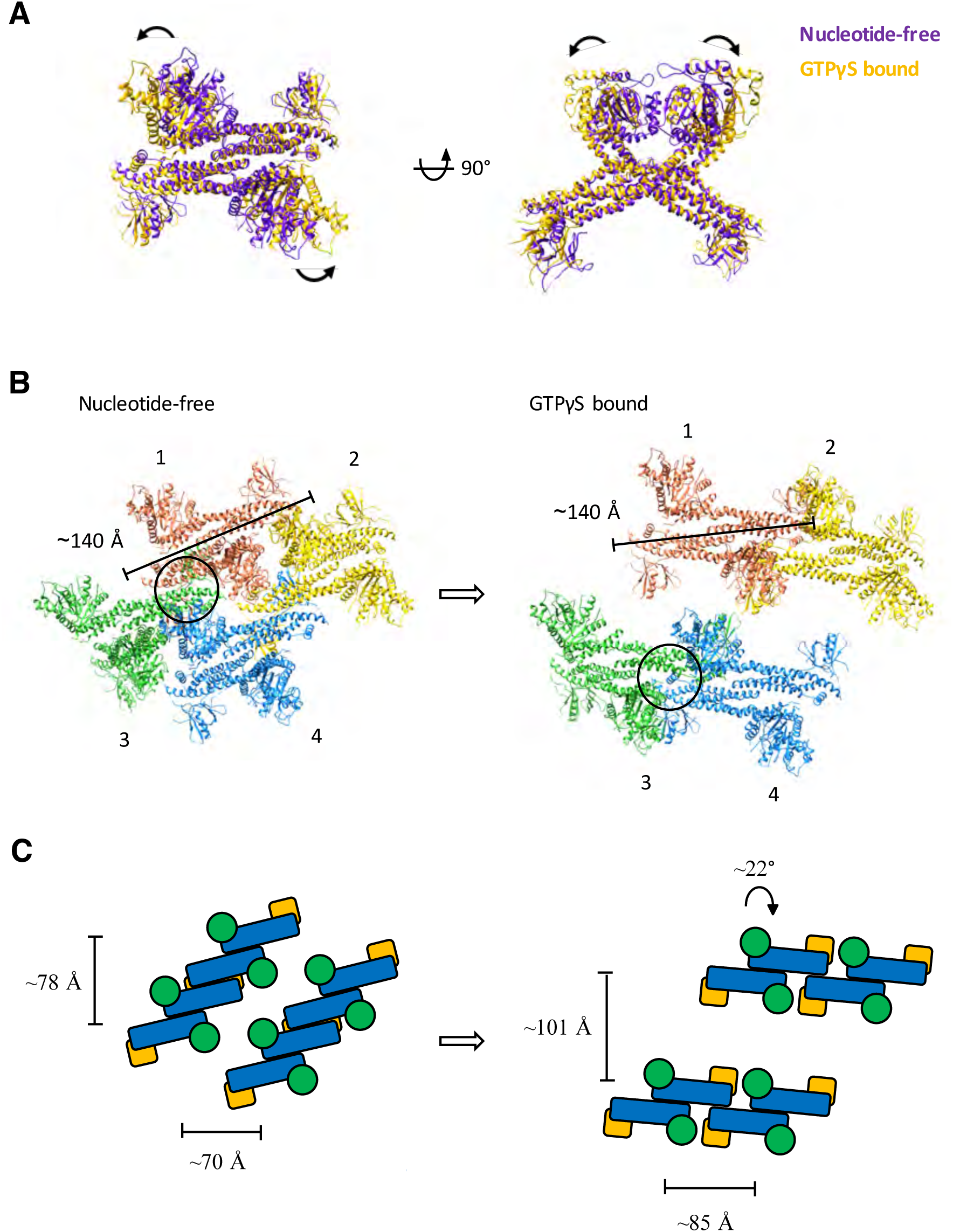
A possible conformational change of S-OPA1 on lipid after GTPγS binding. (**A**) Conformational change of S-OPA1 short dimer after GTPγS binding. The left panel shows the short dimer in a view perpendicular to the helical axis and the right panel shows the view along the axis. Dimer in nucleotide-free state is colored in purple and dimer in GTPγS bound state is colored in yellow. Black arrows indicate the direction of the conformational change. (**B**) Conformational change of S-OPA1 helical lattice after GTPγS binding. Four short dimers are number from 1 to 4. The distances between the tips of stalks in a single building block are labeled and indicated by black lines. Interactions between building blocks as labeled in black cycle. (**C**) Cartoon representation of displacements and rotations between building blocks during GTPγS binding. Distances between neighboring dimers are labeled and indicated by black lines. The black arrows indicate the rotation of each S-OPA1 short dimer.

### A significant conformational change of S-OPA1 after nucleotide binding

Next we further analyzed the conformational changes of S-OPA1 upon nucleotide binding by comparing the structural models of S-OPA1 coated tubes in nucleotide-free and GTPγS bound states. Superimposing the short dimers of S-OPA1 (extended conformation) in two states reveals a significant motion of G domain (**Figure 7A and Movie S2**), yielding an open conformation of S-OPA1 after GTPγS binding.

The assembly array of S-OPA1 on the tube exhibits even more significant changes after GTPγS binding (**Figures 7B, 7C and Movies S3 and S4**). When overlapping the two helical tubes along the vertical direction, a clockwise rotation (∼20°) of the S-OPA1 short dimer was observed and the distance between the short dimers in the same helical rung is increased ∼ 6 Å. Such rotation eliminates the stalk bundle-to-bundle interactions among different helical rungs that are important for the stabilization of nucleotide-free protein-membrane complex (**Figure 3D**). However, a new interface between the short dimers in the same helical rung forms after the structural rotation, which is mediated via a new kind of stalk bundle-to-bundle interactions and thus stabilizes the structure of GTPγS bound protein-membrane complex. Besides the structural rotation, the movement of the S-OPA1 short dimer was also observed, which yields an increased distance of ∼26 Å between dimers in adjacent rungs (**Figures 6C and 6D**).

Overall, GTPγS binding induces a conformational change of S-OPA1 (majorly the movement of G domain) without changing the general architecture of the tube building block. And this conformational change causes the rotation of the S-OPA1 dimers and enlarges the intervals between neighboring dimers, which eventually results an expansion of the helical tube.

## DISCUSSION

Membrane fission and fusion are very important processes in eukaryotic cells for organelle communication, matter exchange and cargo transportation. Owing to the specific double-membrane structure, the fission and fusion of mitochondria are more complicated than plasma membrane and other organelles. With the fruitful structural information of Mfn1/2 and Drp1/Dnm1, the molecular mechanisms of mitochondrial outer membrane fusion and fission have been extensively investigated (Cao et al., 2017; Frohlich et al., 2013; Kalia et al., 2018; Mears et al., 2011; Qi et al., 2016; Yan et al., 2018). However, with the lack of the structure of OPA1, we know little about the molecular mechanism of mitochondrial inner membrane fusion. Compared to Mfn1/2, the primary structure of OPA1 is more like Dyn1 that is physiologically involved in membrane fission. How a membrane-fission-like protein could play a role in membrane fusion has been a puzzle in the field for a long time. In addition, unlike other membrane fusion/fission proteins, OPA1 has multiple isoforms and needs to be processed from a long membrane anchored form (L-OPA1) to a short soluble form (S-OPA1) for an efficient mitochondrial inner membrane fusion, which suggests a strictly regulated dynamics of mitochondrial inner membrane.

In the present study, we investigated the interactions between S-OPA1 and liposome that comprises the phospholipid composition of mitochondrial inner membrane, and then we utilized cryo-electron microscopy approach to resolve the structures of S-OPA1 coated on membrane in both nucleotide-free and GTPγS bound states. Like other dynamin proteins, S-OPA1 can bind to membrane and induce membrane tubulation by forming a helical array. We found such tubulation process is independent with the GTP hydrolysis activity of S-OPA1. While the GTPase activity of S-OPA1can be significantly enhanced upon binding to membrane.

We found S-OPA1 has a typical architecture of dynamin family and comprises a global G domain, a long stalk region and a globular membrane binding (GMD) domain. Analyzing cryo-EM densities reveal two membrane binding sites of S-OPA1, one is located in its GMD domain and another belongs to the extention portion of its stalk region. We further identified an amphipathic helix located in the GMD domain and found this helix contributes partially to membrane binding but determines the membrane tubulation activity of S-OPA1. Indeed, previous literatures have suggested the relations between diseases and the mutations at the stalk and GMD regions in OPA1 (see **Table S2**).

We deduced the building block of S-OPA1 helical array on the membrane is a dimer, which utilizes the tip-to-tip interaction of stalk regions to form the interface. Such dimer organization is significantly different with other dynamin proteins (Faelber et al., 2011; Ford et al., 2011; Reubold et al., 2015). The bundle-to-bundle interactions among stalk regions play key roles in the stability of S-OPA1 coated liposomal tubes. The G domain dimerization interface, which is important for the GTP hydrolysis activity of dynamin proteins, is not formed in the helical array of nucleotide-free S-OPA1, suggesting the GTP hydrolysis is a late-stage event after membrane remodeling by S-OPA1.

We observed a significant conformational change of S-OPA1 after GTPγS binding, which includes the relative motion between G domain and stalk, the global rotation of stalk region and the re-arrangment of S-OPA1 helical lattice. Here we defined the conformation of S-OPA1 in its nucleotide-free state as a close conformation and the one with GTPγS bound as an open conformation. On the contrary to the conventional dynamin proteins that induce a more crowded packing and a constricted tube with a smaller diameter and higher curvature (ready for fission) after nucleotide binding (Chappie et al., 2011; Mears et al., 2011; Sundborger et al., 2014), the conformational change of S-OPA1 from close to open after GTPγS binding results an expanded tube with an increased diameter and reduced membrane curvature. This implies a different role of S-OPA1 in membrane remodeling in comparison with other dynamin proteins.

There is a hypothesis of a mechanochemical mechanism of protein induced membrane remodeling adopted by Dyn1 (Chappie et al., 2011). The protein utilizes the energy generated from binding and hydrolysis of GTP to achieve a conformational change. Such conformational change would deform the membrane and successfully transform the chemical energy into mechanical changes. Since S-OPA1 shows a similar G domain conformational change after nucleotide binding like Dyn1 adopts, it would probably function through the same mechanism on membrane remodeling. Although the resolution by cryo-electron tomography and sub-tomogram averaging is not high, our observation does imply a closer distance of S-OPA1 G domains between two neighbor S-OPA1 short dimer after GTPγS binding, however such distance is still not close enough for G domains dimerization. Thus, the structure of GTPγS bound S-OPA1 we determined is still not the state ready for GTP hydrolysis. We speculate it is due to the subtle difference between GTPγS and GTP, which yields a barrier for future structural re-organization of S-OPA1 helical array to form dimerization of G domains. Thus, in the physiological procedure, with GTP binding a further motion of G domain and S-OPA1 dimer would occur to enable forming G domain dimerization interface ready for GTP hydrolysis.

Previous studies showed that S-OPA1 alone could not trigger the fusion of mitochondrial inner membrane and S-OPA1 needs to work together with L-OPA1 (Ban et al., 2017; Del Dotto et al., 2017; Song et al., 2007). During mitochondrial inner membrane fusion, L-OPA1 would perform a more important role while the addition of S-OPA1 would significantly increase the fusion efficiency (Ban et al., 2017). In addition, the GTPase activity of S-OPA1 is indispensable for its fusion related function (Ban et al., 2017). Those reveal an assumption that S-OPA1 will assist the function of L-OPA1 with its GTPase activity during mitochondrial inner membrane fusion. Thus, we speculate S-OPA1 facilitates the membrane fusion by oligomerization together with L-OPA1, helping with the GTP hydrolysis and conformational change of L-OPA1. Studies of Mgm1, the homolog of OPA1 in yeast, indicated a similar situation in which the GTPase activity of L-Mgm1 (long isoform) is inhibited because of the restriction of G domain movement by the transmembrane helix (DeVay et al., 2009). The S-Mgm1 (short isoform) would then bind to facilitate the generation of G dimer and accelerate GTP hydrolysis.

Though our current studies tried to understand OPA1 induced mitochondrial inner membrane fusion, the precise cooperating mechanism of L-OPA1 and S-OPA1 remains further elucidated. The high-resolution information of S-OPA1 and L-OPA1 in different states need determined to unravel the molecular mechanism of mitochondrial inner membrane fusion.

## MATERIALS AND METHODS

### Protein expression and purification

cDNA corresponding to the short S1 isoform of OPA1 was sub-cloned into the pET32M-3C expression vector (from Wei Feng’s Lab, IBP, CAS) with a N-terminal Trx tag and a followed His6 tag. And there is a prescission protease cleavage site exist between His tag and protein sequence. Mutants were constructed in pET32M-3C/OPA1-S1 via PCR. All proteins were expressed in Transetta (DE3) bacteria cells (Transgene) and purified under the following procedure. Cultures were grown at 37 °C until OD 600 nm reaching 0.8 and induced with 0.2 mM IPTG for 18 hr at 16°C. The cells were collected by centrifugation. Bacteria pellets were resuspended in lysis buffer containing 20 mM Tris-HCl (pH 8.0), 150 mM NaCl and 1 tablet of protease inhibitors cocktail (Roche) and disrupted with ultra-sonication. Lysates were incubated with Ni-NTA beads (Roche). After washing with the buffer containing 20 mM Tris-HCl (pH 8.0), 150 mM NaCl, 1mM DTT, and 10 mM imidazole, protein was cleaved by prescission protease at 4°C overnight. After cleavage, protein was eluted with 20 mM Tris-HCl (pH 8.0), 150 mM NaCl, 1 mM DTT, 20 mM imidazole. The eluted protein fraction was further purified by gel filtration chromatography using a Superdex 200 10/300 GL column (GE Healthcare) in the buffer of 20 mM Tris-HCl (pH 8.0), 150 mM NaCl, and 1 mM DTT. The elution volume of the column was pre-calibrated using standard protein molecular weight markers. Purified proteins were frozen in liquid nitrogen and stored at −80 °C.

### Preparation of S-OPA1 coated tubes

The lipids (Avanti Polar Lipids) were mixed in the following ratio: 45% palmitoyl-2-oleoyl-sn-glycero-3-phosphocholine (POPC), 22% 1-palmitoyl-2-oleoyl-sn-glycero-3-phosphoethanolamine (POPE), 8% L-α-lysophosphatidylinositol (PI), and 25% 1’,3’-bis[1,2-dioleoyl-sn-glycero-3-phospho]-sn-glycerol (cardiolipin). The indicated ratios of lipids were mixed in a chloroform solution, evaporated for 4 hr in a vacuum desiccator and rehydrated in the buffer of 20 mM Tris–HCl (pH 8.0), 1 mM EGTA, and 1 mM MgCl_2_ to a final concentration of 4 mg/ml. The resulting multi-lamellar liposomes were put through five freeze/thaw cycles to make unilamellar liposomes. S-OPA1 protein was then mixed with unilamellar liposomes 1:1 (m:m) at a final concentration of 1mg/ml in the buffer containing 20 mM HEPES (pH 8.0), 1 mM EGTA, and 1 mM MgCl_2_. The mixture was incubated at 16 °C for 30 min before preparing cryo-EM samples. For tubes incubate with GTP and GTP non-hydrolyzed analogous, 10 mM nucleotide was then added with a final concentration of 1mM, and the mixture was incubated for another 30 min.

For negative staining EM, 5 μl of protein-lipid tubes was applied to glow-discharged continuous carbon films and stained with uranyl acetate (2% w/v) for 1 min. Samples were visualized using a Tecnai Spirit electron microscope (ThermoFisher Scientific) operating at 120 kV and image were recorded with an Eagle camera.

Cryo-EM grids were prepared with Vitrobot Mark IV (ThermoFisher Scientific) under 100% humidity. 3 μl of protein-lipid tubes was applied to glow-discharged Quantifoil R2/1 holy carbon grids, blotted, and plunged into liquid ethane. For grids using for tomography data collection, homemade protein A coated colloidal gold was added as a fiducial marker.

### Cryo-electron microscopy

Images for helical reconstruction were recorded on a cryo-electron microscope Titan Krios (ThermoFisher Scientific) operating at 300kV using SerialEM software (Mastronarde, 2005). A Falcon-IIIEC camera (ThermoFisher Scientific) was used at a calibrated pixel size of 1.42Å. A combined total dose of 50 e/Å^2^ was applied with each exposure. Images were collected at 2-4 μm underfocus.

Tilt series data were collected on a cryo-electron microscope Titan Krios G2 (ThermoFisher Scientific) using SerialEM software (Mastronarde, 2005), with a K2 direct electron detector (Gatan) operating in counting mode. Tilt series data were typically collected from ±45° with 3° tilt increments at 3-5 μm underfocus. A combined dose of about 90 e/Å^2^ was applied over the entire series.

### Helical reconstruction

In total, 2112 movie stacks were collected. Motion correction and defocus estimation for all these micrographs were performed using MotionCorr2 (Zheng et al., 2017) and GCTF (Zhang, 2016) respectively. Micrographs with ice contamination, poor Thon rings, too large defocus values (greater than 3 μm) were excluded before filament boxing. Good micrographs were then multiplied by their theoretical contrast transfer function (CTF) for initial correction of CTF. 511 S-OPA1 tubes were boxed using e2helixboxer.py in the package of EMAN2 (Tang et al., 2007) with a 480 px box width. An initial segment was stack generated from all these tubes with an overlap of 90% and an initial 3D model was generated by back projection method using these segments and assigning random azimuthal angles to them. The initial 3D model was then interpolated into different ones with various diameters, and diameter classification was performed through supervised 2D classification where models were generated by projecting the 3D models with various diameters. Then for each diameter class, the diffraction pattern for each tube was calculated and further classified. A main class of tubes at ∼ 53 nm diameter was sorted corresponding to its diameter and diffraction pattern, and contains 6644 segments. The segment stack of the selected class was then regenerated with the box size of 480 px and the box overlap of ∼ 94%. Initial helical parameters were calculated by indexing the layer lines in the power spectrum of the boxed tubes. An initial helical rise of 27.0Å and twist of 20.87° were obtained and used for helical reconstruction through a real space helical reconstruction algorithm IHRSR (Egelman, 2000, 2007). The helical parameters finally converged to 25.87 Å for the helical rise, and 20.86° for the helical twist. Then summed CTF^2^ was divided for the final CTF correction of the map reconstructed by IHRSR. SPIDER (Shaikh et al., 2008) was used for negative B-factor sharpening. Resolution of the final map was estimated based on the gold standard Fourier shell correlation (FSC)0.143 criterion.

### Tomographic reconstruction and sub-volume averaging

Fiducial marker based tilt series alignment and gold erasure were performed using AuTom (Han et al., 2017). And the tomographic reconstructions were performed using IMOD (Kremer et al., 1996) with 2 times binning. No CTF correction was performed at this step. For tomographic reconstruction, the radial filter options were set at 0.35 cut off and 0.05 fall off. The sub volumes picked in IMOD were extracted by RELION 1.4 (Scheres, 2012) and CTF model of each particles was generated through RELION script that called CTFFIND4 (Rohou and Grigorieff, 2015). Then 3D classification was carried with CTF correction and the particles from selected classes were used for the final refinement. The initial model for 3D classification and refinement is the random averaging of all particles and low-pass filtered to 60 Å. Reported resolutions are based on the gold-standard Fourier shell correlation (FSC) 0.143 criterion.

### Sedimentation assay

Sedimentation assay was carried on as previous study (Ban et al., 2010). Proteins was diluted to a concentration of 0.2 mg/ml in 20 mM Tris-HCl (pH 8.0), 300 mM NaCl, 1mM MgCl_2_, 1mM EGTA, 1 mM DTT. Liposomes are the same as the above tubulation assay. The liposomes were directly added to the protein solution at a final concentration of 0.2 mg/ml and incubated at room temperature for 30 min. Samples were centrifuged at 250,000 g in a S140AT rotor (Hitachi) for 20 min at 4°C. The supernatant and pellet were analyzed by SDS–PAGE.

### GTPase activity assay

GTPase reactions were performed as previous studies (Ban et al., 2010) with 0.1 mg/ml protein and 0.1 mg/ml liposomes in 20 mM HEPES (pH 7.5), 1 mM EGTA, and 1mM MgCl_2_. GTP hydrolysis was quantified by monitoring the free phosphate concentration using a malachite green assay (Leonard et al., 2005). Reactions were initiated by the addition of GTP to 100 mM and incubation at 37°C. 20 μl mixtures were quenched with 5 μl of 0.5 M EDTA at regular time points. After the addition of 150 μl malachite green solution, the free phosphate concentration was monitored by the absorbance at 650 nm in a 96-well plate reader (EnSpire 2300).

Calculation of Km and Kcat is based on Lineweaver-Burk plot with the measurement at 6 different GTP concentration. The final GTP concentration was set to 0.4, 0.5, 0.6, 0.8, 1.0, and 1.4 mM individually. Reactions were initiated by the addition of 10 μl GTP solution to a 10 μl protein or protein liposome mixtures in 20 mM HEPES (pH 7.5), 1 mM EGTA, and 1mM MgCl2. After 1∼2 h incubation, which was set based on the activity of different mutants, at 37 °C, the reactions were quenched with 10 μl 0.1 M EDTA. The phosphate was quantified using malachite green assay. 150 μl malachite green solution was added and the absorbance at 650 nm is monitored in a 96-well plate reader (EnSpire 2300). All experiments were repeated at least three times.

## DATA AVAILABILITY

Cryo-EM maps and models of S-OPA1 coated liposomal tubes have been deposited into Electron Microscopy Data Bank and Protein Data Bank with the accession codes EMD-XXXX/PDB-XXXX for the helical reconstruction of nucleotide-free state, EMD-XXXX for the tomographic reconstruction of nucleotide-free state and EMD-XXXX/PDB-XXXX for the tomographic reconstruction of GTPγS bound state, respectively.

## AUTHOR CONTRIBTUIONS

F. S. initiated and supervised the project. D.Z. performed all experiments and collected all data. Y.Z. performed image processing of helical reconstruction. D.Z. and T.N. performed image processing of cryo-electron tomography and sub-tomogram averaging. J.M. cloned the gene and performed initial protein expression. W.C. performed initial cryo-EM work of S-OPA1 coated tube. X.P. helped on liposome preparation and co-sedimentation experiments. Y.Z. helped on protein production and purification. D.Z., Y.Z., and F.S. analyzed the data and wrote the manuscript.

## ACKNOWLEDGMENTS

We would like to thank Prof. Edward H. Egelman from University of Virginia for his generous help on image processing of helical reconstruction. We are grateful to Xue Wang (F.S. lab) for her initial trials of tubulation experiments. We would also like to thank Ping Shan and Ruigang Su (F.S. lab) for their assistances. This work was supported by National Natural Science Foundation of China (31770794), grants from Chinese Academy of Sciences (XDB08030202) and the Ministry of Science and Technology of China (2017YFA0504700). All the EM works were performed at Center for Biological Imaging (CBI, http://cbi.ibp.ac.cn), Institute of Biophysics, Chinese Academy of Sciences.

## COMPLIANCE WITH ETHICAL STANDARDS

All authors declare that they have no conflict of interest. All institutional and national guidelines for the care and use of laboratory animals were followed.

