## Supplementary Figures and Tables for "Cryo-EM structures reveal interactions of S-OPA1 with membrane and changes upon nucleotide binding"

### Supplementary Methods

#### Size exclusion chromatography coupled with multi angle light scattering

The size of S-OPA1 in solution is determined by static multiangle light scattering (MALS) coupled with gel filtration. The size-exclusion chromatography column (PROTEIN KW-803, Shodex) is equilibrated with 20 mM Tris (pH 8.0), 150 mM NaCl, and 100 μl of 1mg/ml purified S-OPA1 was applied. The detector DAWN HELEOS II (Wyatt) was used to measure the mass distribution. Data were analyzed using the provided ASTRA software.

#### Chemical crosslink assay

S-OPA1 and Δ196-252 was diluted to 0.3 mg/ml in 50 mM HEPES (pH 7.4), 300 mM NaCl and 1 mM DTT. The amine-reactive crosslinker bis(sulfosuccinimidyl) suberate (BS3; Thermo Fisher Scientific) was added to a final concentration of 50 μM. After a 15 min incubation at room temperature, the crosslinking reaction was quenched with 50 mM Tris (pH 8.0). Crosslinked products were analyzed using a 3–8% Tris–acetate PAGE.

### Legends for Supplementary Materials

**Figure S1. Characterization of S-OPA1 and helical image processing of S-OPA1 coated tubes at the nucleotide-free state.** (**A**) SEC-MALS (size exclusion chromatography coupled with multi angle light scattering) profile of full-length S-OPA1 (WT) and the molecular weight of the oligomer was determined. (**B**) Chemical crosslinking (using BS3) SDS-PAGE of the full-length S-OPA1 and the truncation form △196-252. (**C**) Different diffraction patterns of the helical tubes. (**D**) Diameter distribution of the helical tubes. (**E**) Diffraction patterns of the class averaged tube (left) and projection of the reconstructed map (right). (**F**) Indexing of the layer lines of S-OPA1 coated tube. (**G**) Structure prediction of S-OPA1 using Phyre2 and I-TASSER. Related to **Figures 1 and 2**.

**Figure S2. Sub-tomogram averaging of full length and truncated S-OPA1 coated tubes at nucleotide-free state.** (**A**) The 3D averaged map of truncated S-OPA1 coated tube viewed at different slicing positions in **Figure 2B**. The bottom panel shows the cross section at the radical direction. The map is colored with the same scheme in **Figure 2**. The dashed lines denote the slicing positions in the above panels. The structural model derived from helical reconstruction (Figure 3) was directly superimposed into the map and shown on the right panel. (**B**) The 3D averaged map of full length S-OPA1 coated tube viewed the same as (A). The structural model derived from helical reconstruction (Figure 3) was directly superimposed into the map and shown on the right panel. Related to **Figure 2 and 3**.

**Figure S3. Docking crystallized fragments of dynamin1 into cryo-EM map of nucleotide-free S-OPA1 coated tube.** (**A**) Docking of the crystal structure of dimerized G domain ($GG_{GDP.AlF4^-}$, PDB ID 2X2E) into the head region of the map. Left, sideview; Right, end-on view. (**B**) 2D radical cross section of the map. The two membrane interacting sites of S-OPA1 are indicated with the black arrows. (**C**)

Two possible building blocks of S-OPA1 coated tube at nucleotide-free state. Short dimer is in green and long is dimer in pink. Related to **Figure 3**.

**Figure S4. S-OPA1 mutants on G domain.** Sequence alignment of partial G domains among different dynamin proteins. Black arrows indicate the mutation sites. Related to **Figure 4**.

**Figure S5. A predicted amphipathic helix of S-OPA1.** (**A**) Sequence alignment of S-OPA1 and dynamin1 at their PH/GMB domains. The region of dynamin 1 PH domain is labeled. (**B**) Helical wheel analysis of the amphipathic helix. Related to **Figure 5**.

**Figure S6. Cryo-electron tomographic image processing of S-OPA1 coated tubes after adding GTPγS.** (**A**) Cryo-EM images of full length S-OPA1 (WT) and truncated S-OPA1 (Δ196-252) coated tubes at nucleotide-free state and after adding GTP, GMPPCP, GMPPNP or GTPγS. Scale bar, 100 nm. (**B**) Diameter distribution of the expanded S-OPA1 tubes after adding GTPγS. (**C**) One selected tilt micrograph of tomographic dataset with the tilt angle of 0°. (**D**) Flow chart of tomography and sub-tomogram averaging image processing procedure. (**E**) Angular distribution of sub-tomograms used for final reconstruction. (**F**) The gold standard FSC (Fourier Shell Correlation) curve and the estimated resolution of the sub-tomogram averaged map at the threshold of 0.143. Related to **Figure 6**.

**Figure S7. Docking crystallized fragments of dynamin1 into cryo-EM map of S-OPA1 coated tube at GTPγS bound state.** (**A**) Docking of the crystal structure of dimerized G domain ($GG_{GDP.AlF4}$-, PDB ID 2X2E) into the head region of the map. Left, sideview; Right, end-on view. (**B**) Two possible building blocks of S-OPA1 coated tube at GTPγS bound state. Short dimer is in green and long is dimer in pink. Related to **Figure 6**.

**Figure S8. The rational analysis of S-OPA1 long dimer and comparison of S-OPA1 short dimer with other dynamin protein dimers.** (**A**) The invalidity of S-OPA1 long dimer is proved by analyzing the conformational change of S-OPA1 helical lattice after adding GTPγS. (**B**) Comparing of S-OPA1 short dimer with other dynamin protein dimers including Dyn1 (PDB ID 3SNH), MxA (PDB ID 3SZR) and Drp1 (PDB ID 4BEJ). The black circles highlight the conventional interface in dynamin dimer. Related to **Figure 7**.

**Table S1. Enzymatic Km and Kcat of S-OPA1 and its mutants.** The data presented come from three independent experiments (**Data S1**).

**Table S2. Disease related mutants of S-OPA1. Data was obtained from UniProt (https://www.uniprot.org).**

**Data S1. Original data of enzymatic assays of S-OPA1 and its mutants and and detailed data processing.**

**Movie S1. Structure of S-OPA1 coated tube in a nucleotide-free state.**

**Movie S2. Conformational changes of S-OPA1 monomer and its short dimer after GTPγS binding.**

**Movie S3. Conformational change between S-OPA1 helical lattice after GTPγS binding.**

### Figure S1

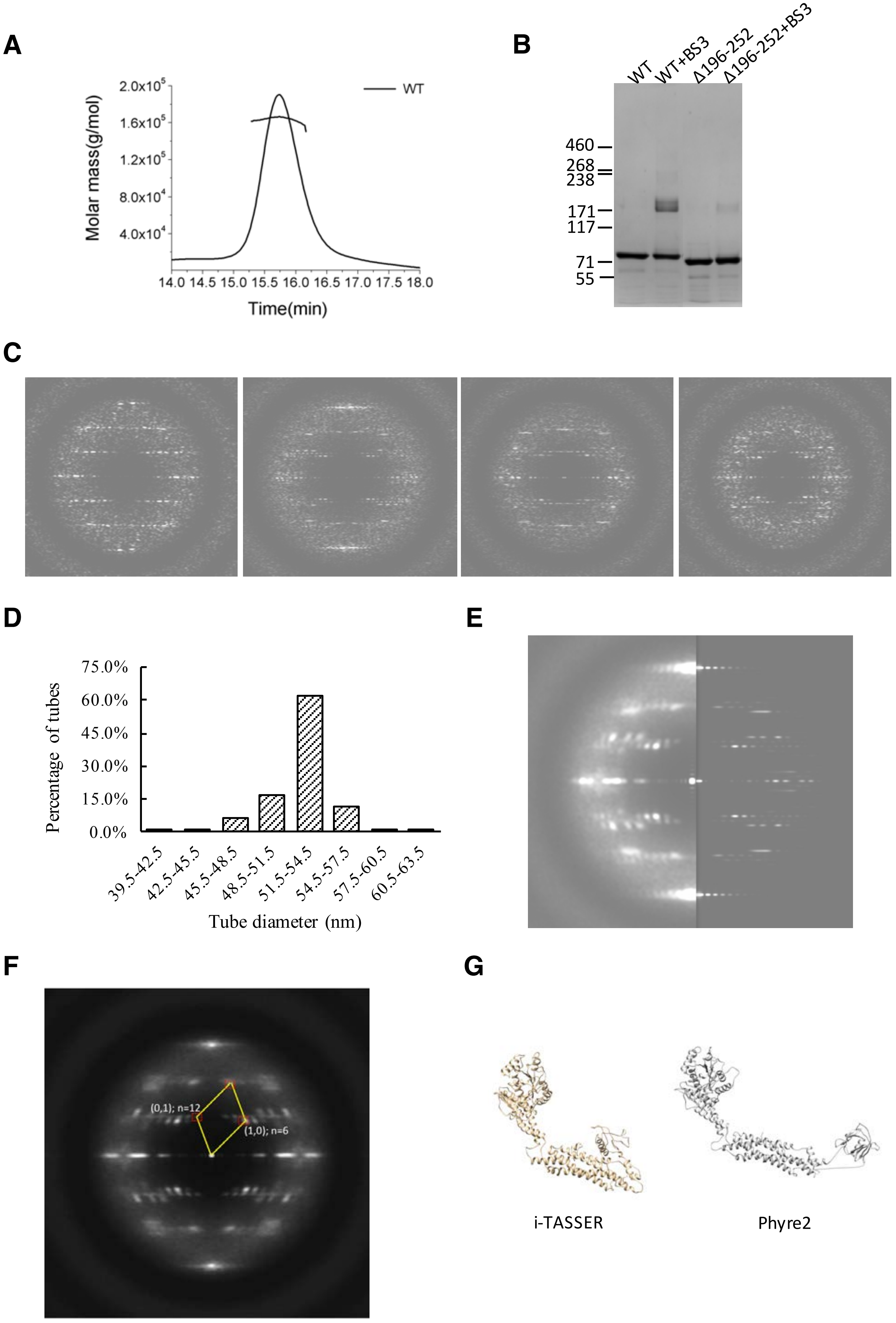
A
B
C
D
E
F
G
WT
Molar mass(g/mol)
2.0x10^5
1.6x10^5
1.2x10^5
8.0x10^4
4.0x10^4
14.0 14.5 15.0 15.5 16.0 16.5 17.0 17.5 18.0
Time(min)
WT
WT+BS3
Δ196-252
Δ196-252+BS3
460
268
238
171
117
71
55
Percentage of tubes
75.0%
60.0%
45.0%
30.0%
15.0%
0.0%
39.5-42.5
42.5-45.5
45.5-48.5
48.5-51.5
51.5-54.5
54.5-57.5
57.5-60.5
60.5-63.5
Tube diameter (nm)
(0,1); n=12
(1,0); n=6
i-TASSER
Phyre2

### Figure S2

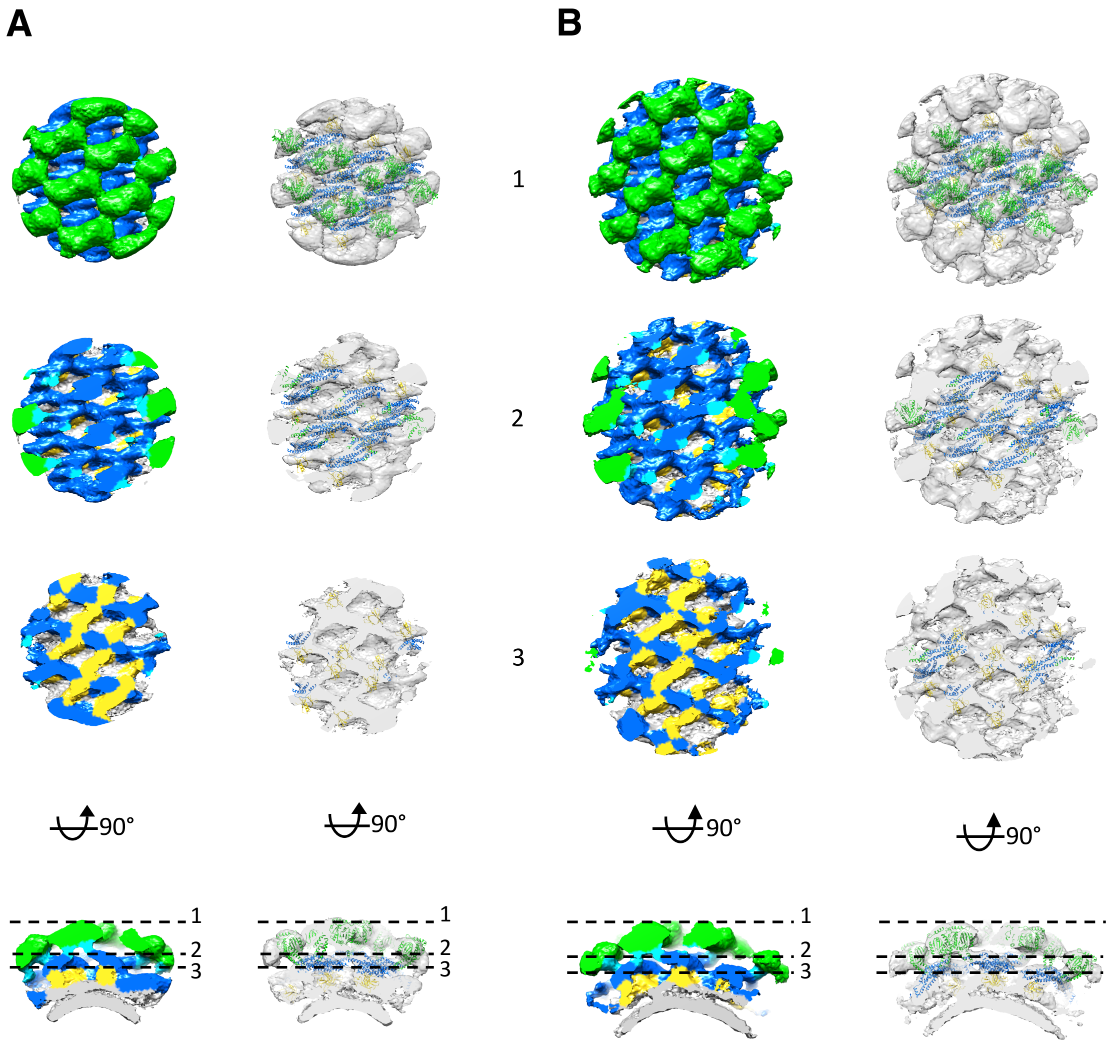

**Figure S3**

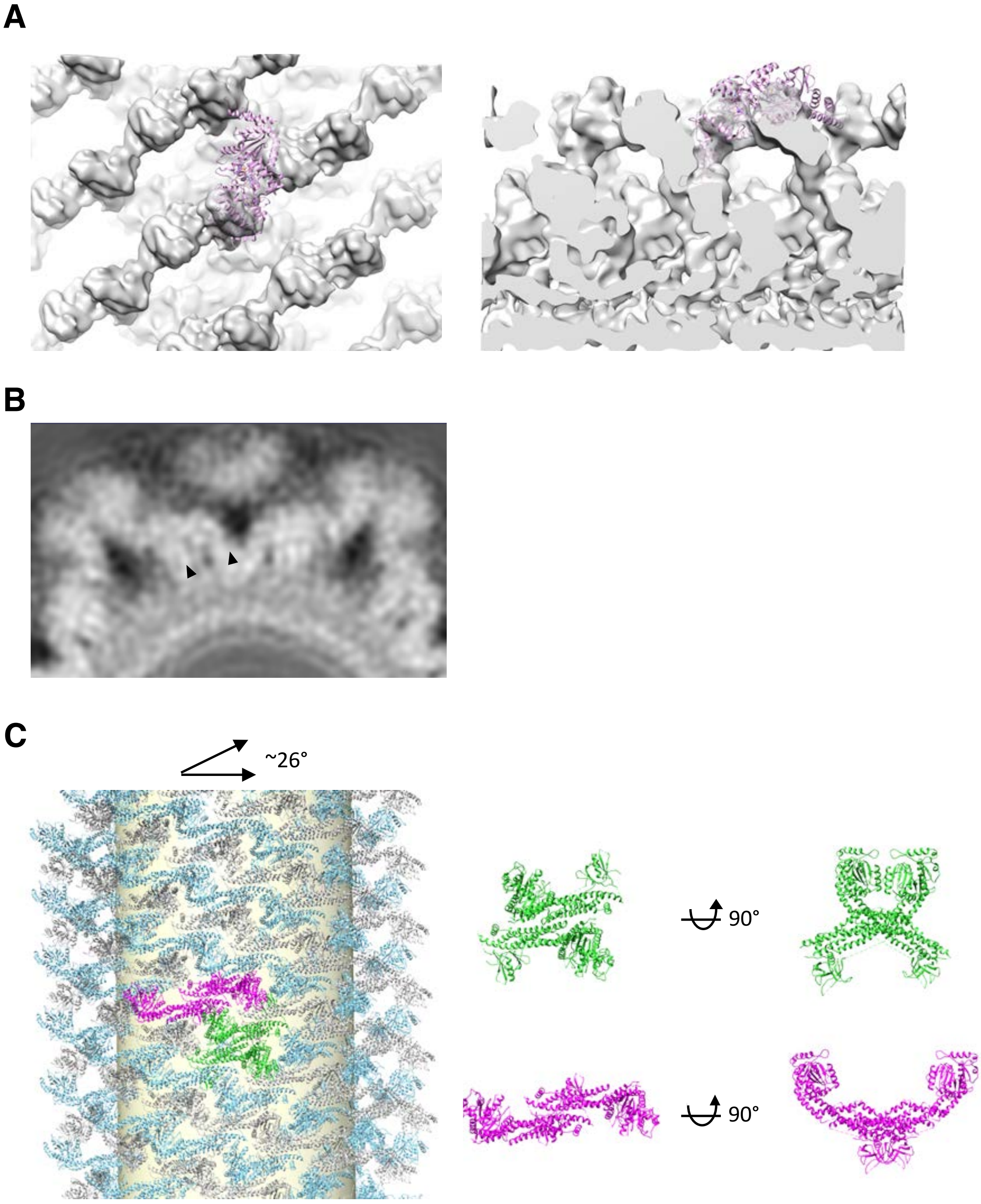

**Figure S4.**

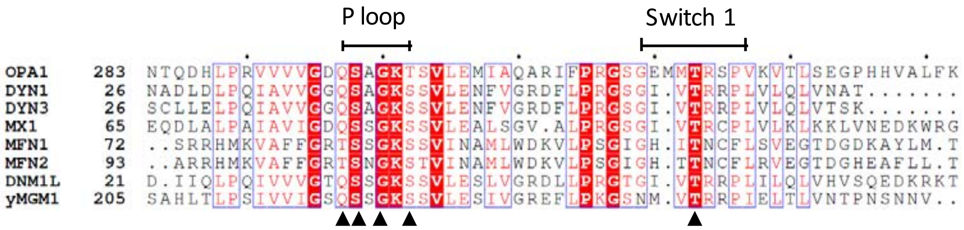
P loop
Switch 1
OPA1 283 NTQDHLPRVVVVGDQSAGKTSVLEMIAQARIFPRGSGEMMTRSPVKVTLSEGPHHVALFK
DYN1 26 NADLDLPQIAVVGGQSAGKSSVLENFVGRDFLPRGSGI.VTRRPLVLQLVNAT.......
DYN3 26 SCLLELPQIAVVGGQSAGKSSVLENFVGRDFLPRGSGI.VTRRPLVLQLVTSK.......
MX1 65 EQDLALPAIAVIGDQSSGKSSVLEALSGV.ALPRGSGI.VTRCPLVLKLKKLVNEDKWRG
MFN1 72 ..SRRHMKVAFFGRTSSGKSSVINAMLWDKVLPSGIGH.ITNCFLSVEGTDGDKAYLM.T
MFN2 93 ..ARRHMKVAFFGRTSNGKSTVINAMLWDKVLPSGIGH.TTNCFLRVEGTDGHEAFLL.T
DNM1L 21 D.IIQLPQIVVVGTQSSGKSSVLESLVGRDLLPRGTGI.VTRRPLILQLVHVSQEDKRKT
yMGM1 205 SAHLTLPSIVVIGSQSSGKSSVLESIVGREFLPKGSNM.VTRRPIELTLVNTPNSNNV..

### Figure S5

**A**

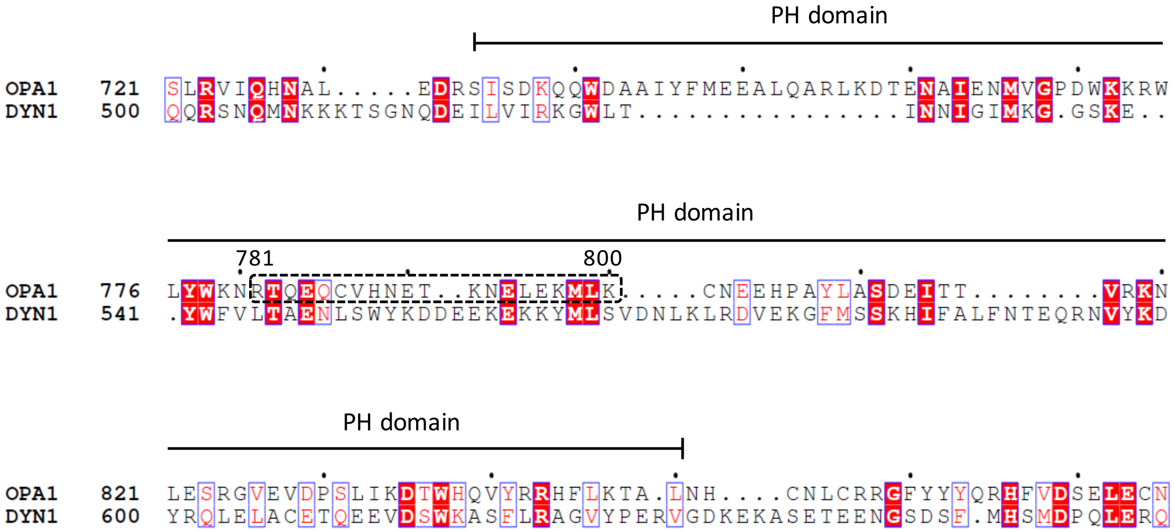


**B**

794-800 ELEKMLK

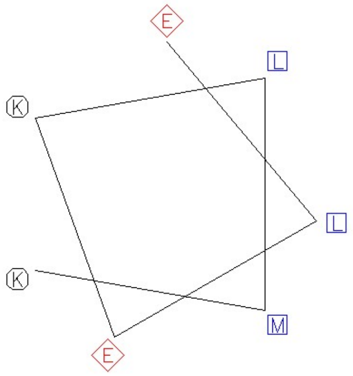

### Figure S6

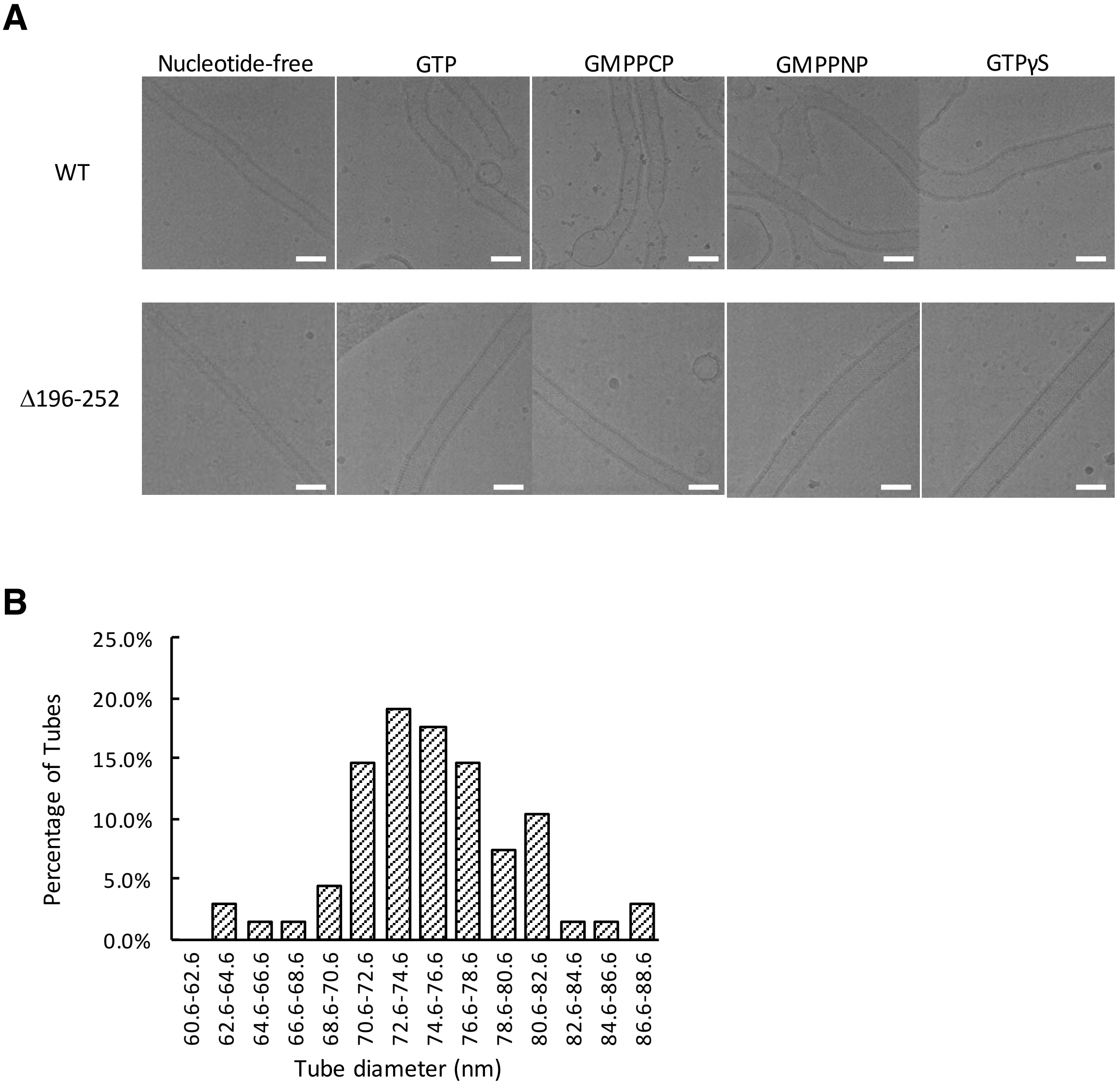
A
Nucleotide-free
GTP
GMPPCP
GMPPNP
GTPγS
WT
Δ196-252
B
25.0%
20.0%
15.0%
10.0%
5.0%
0.0%
Percentage of Tubes
60.6-62.6
62.6-64.6
64.6-66.6
66.6-68.6
68.6-70.6
70.6-72.6
72.6-74.6
74.6-76.6
76.6-78.6
78.6-80.6
80.6-82.6
82.6-84.6
84.6-86.6
86.6-88.6
Tube diameter (nm)

C

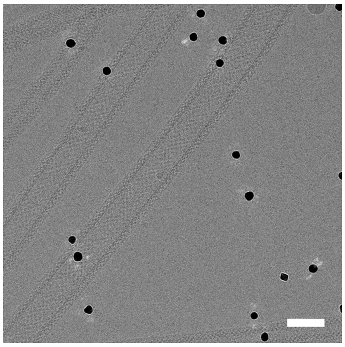

D

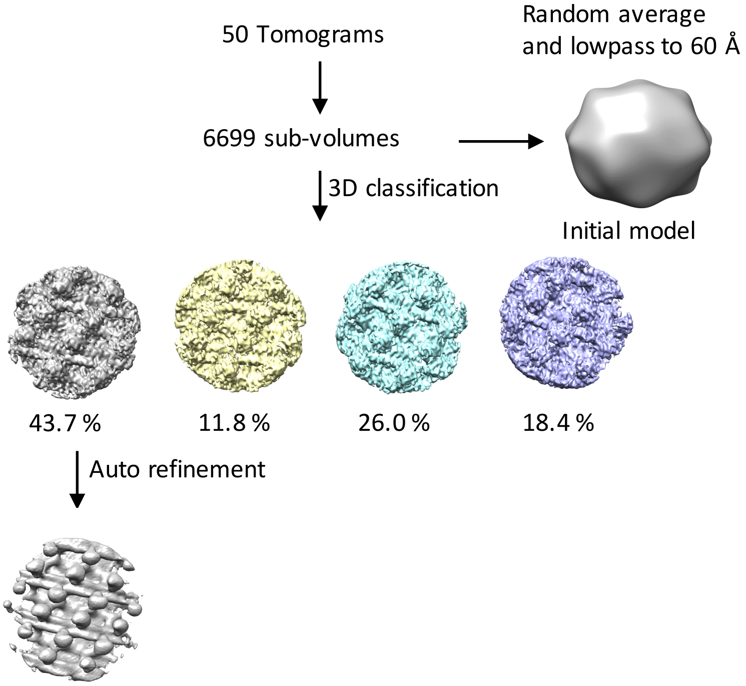

50 Tomograms
Random average
and lowpass to 60 Å
6699 sub-volumes
3D classification
Initial model
43.7 %
11.8 %
26.0 %
18.4 %
Auto refinement


E

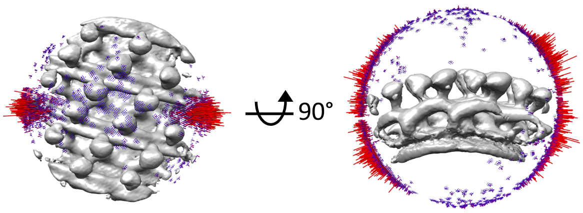

90°


F

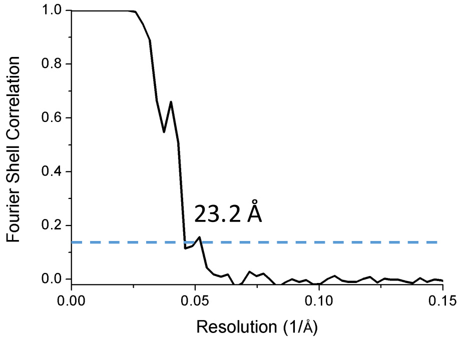

Fourier Shell Correlation
Resolution (1/Å)
23.2 Å
1.0
0.8
0.6
0.4
0.2
0.0
0.00
0.05
0.10
0.15

#### Figure S7

**A**

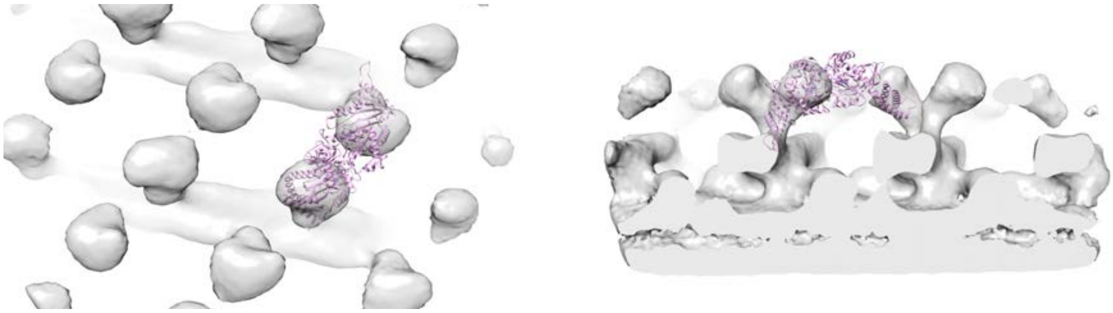

**B**

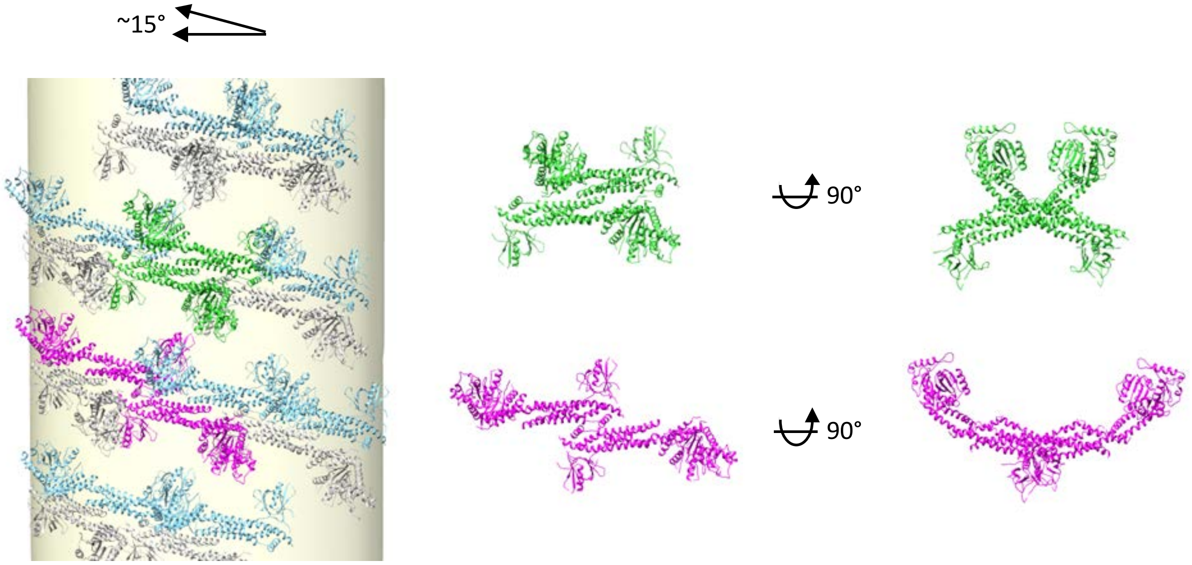

### Figure S8

**A**

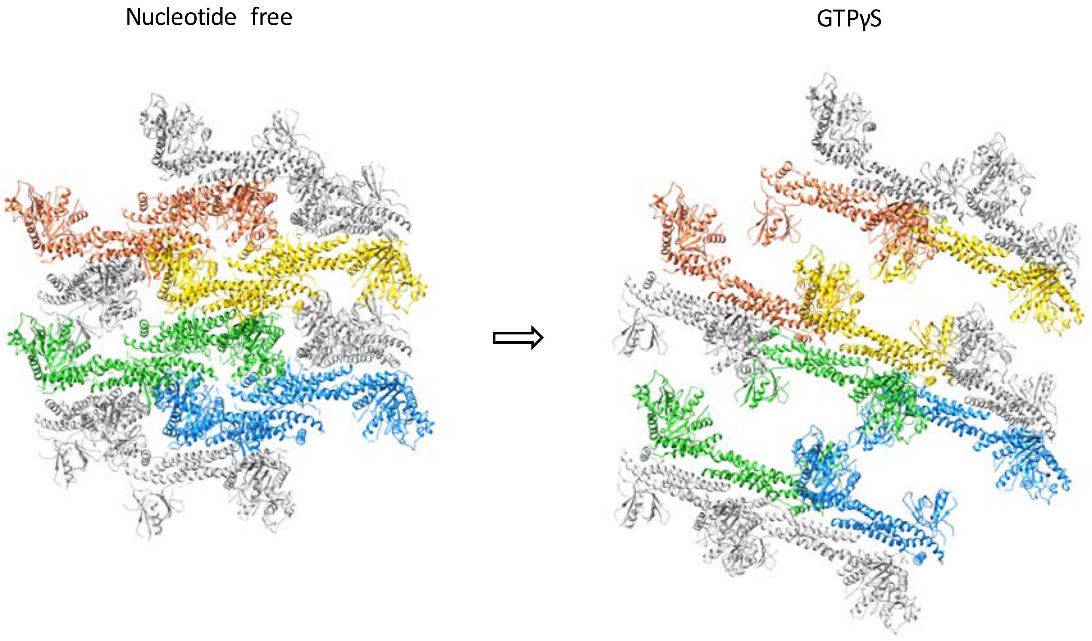

Nucleotide free
GTPγS


**B**

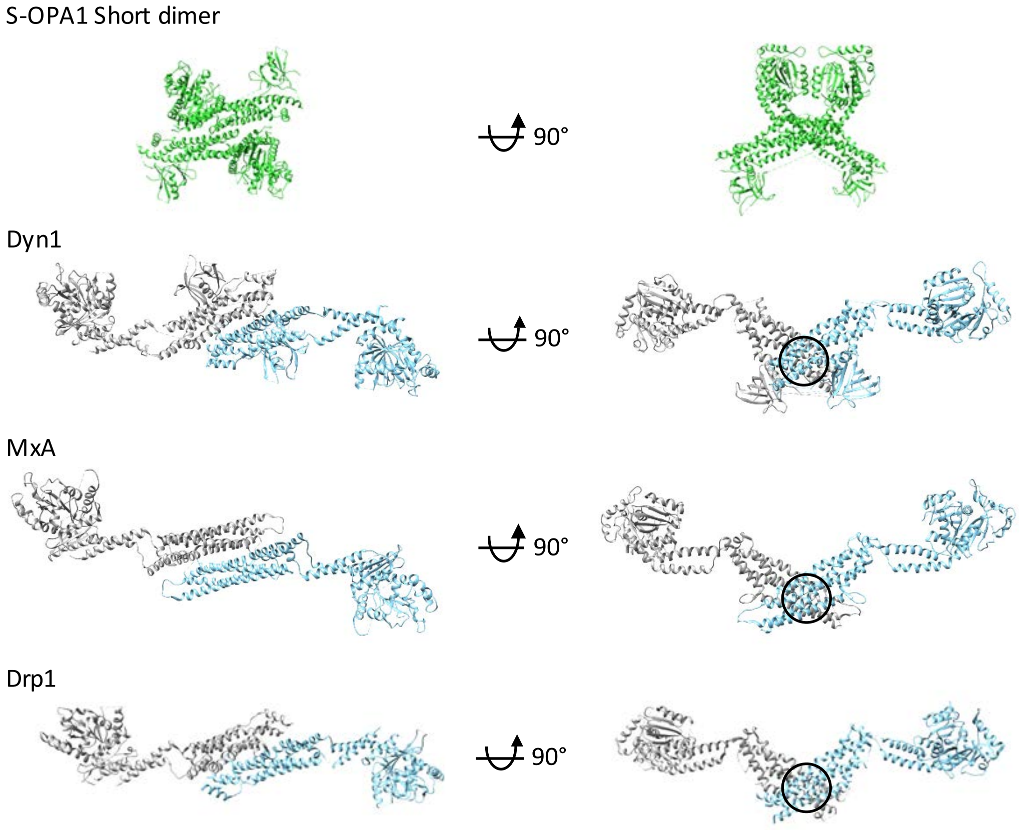

S-OPA1 Short dimer
90°
Dyn1
90°
MxA
90°
Drp1
90°

**Table S1. Enzymatic Km and Kcat of S-OPA1 and its mutants. The data presented come from three independent experiments (Data S1).**

| | No liposome | | | | With liposome | | | |
|---|---|---|---|---|---|---|---|---|
| | Km (mM) | Stdev | Kcat (1/min) | Stdev | Km (mM) | Stdev | Kcat (1/min) | Stdev |
| WT | 1.2 | 0.7 | 3.9E-04 | 1.5E-04 | 5.0 | 5.5 | 2.3E-02 | 2.2E-02 |
| Δ196-252 | 0.4 | 0.2 | 7.2E-05 | 1.8E-05 | 0.8 | 0.6 | 1.1E-02 | 4.5E-03 |
| Q297E | 0.5 | 0.2 | 1.2E-04 | 1.8E-05 | 9.4 | 14.1 | 2.1E-03 | 3.0E-03 |
| S298A | 0.4 | 0.1 | 1.2E-04 | 1.2E-05 | 0.6 | 0.3 | 2.1E-04 | 5.6E-05 |
| G300E | 0.9 | 0.4 | 1.5E-04 | 3.9E-05 | 0.6 | 0.5 | 1.5E-04 | 5.7E-05 |
| T302N | 0.7 | 0.1 | 2.2E-04 | 1.7E-05 | 1.1 | 0.3 | 4.2E-04 | 6.6E-05 |
| T323A | 0.9 | 0.2 | 1.8E-04 | 2.6E-05 | 2.4 | 1.1 | 7.8E-04 | 2.8E-04 |
| 794-800A | 0.6 | 0.2 | 2.2E-04 | 3.0E-05 | 1.1 | 0.3 | 9.1E-04 | 1.6E-04 |
| E794AE796A | 0.5 | 0.1 | 5.3E-04 | 4.1E-05 | 2.8 | 10.4 | 5.3E-03 | 1.5E-02 |
| K797AK800A | 0.5 | 0.5 | 2.2E-04 | 9.1E-05 | 1.5 | 0.6 | 9.4E-03 | 2.7E-03 |
| L795EM798EL799E | 1.9 | 1.1 | 3.8E-04 | 1.7E-04 | 1.1 | 0.6 | 3.9E-04 | 1.4E-04 |
| L795AM798AL799A | 0.9 | 0.2 | 3.6E-04 | 4.1E-05 | 1.3 | 0.2 | 1.1E-03 | 1.4E-04 |

**Table S2. Disease related mutants in S-OPA1. Data was obtained from UniProt (https://www.uniprot.org).**

| Optic atrophy 1 (OPA1) | | | | |
|---|---|---|---|---|
| Position(s) | Mutant | Domain (Predicted) | Description | Reference |
| 8 | A → S | G domain | Unknown pathological significance. | (1) |
| 38 – 43 | Missing | G domain | | (2) |
| 80 | Y → C | G domain | | (1) |
| 95 | T → M | G domain | | (3) |
| 102 | Y → C | G domain | | (3) |
| 270 | E → K | G domain | | (4) |
| 272 | L → P | G domain | | (5) |
| 273 | D → A | G domain | | (4) |
| 290 | R → Q | G domain | | (4, 6-8) |
| 290 | R → W | G domain | | (4) |
| 293 – 294 | Missing | G domain | | (3) |
| 300 | G → E | G domain | Loss of GTPase activity; loss of function in promoting mitochondrial fusion. | (8-10) |
| 310 | Q → R | G domain | | (3) |
| 324 – 326 | Missing | G domain | | (11) |
| 330 | T → S | G domain | | (12) |
| 357 | A → T | G domain | In DOA+ and OPA1. | (3, 13) |
| 377 | V → I | G domain | | (12) |
| 382 | I → M | G domain | In OPA1 and BEHRS. | (3, 14, 15) |
| 384 | L → F | G domain | | (8) |
| 396 | L → P | G domain | | (3) |
| 396 | L → R | G domain | | (2) |
| 400 | P → A | G domain | | (16) |
| 429 – 430 | Missing | G domain | | (3) |
| 430 | N → D | G domain | | (3) |
| 432 | Missing | G domain | | (2, 7) |
| 438 | D → V | G domain | | (4) |
| 439 | G → V | G domain | In DOA+ and OPA1; decreased GTPase activity; loss of function in promoting mitochondrial fusion. | (10, 13, 17) |
| 445 | R → H | G domain | In DOA+ and OPA1; decreased GTPase activity; loss of function in promoting mitochondrial fusion. | (10, 13, 18-20) |
| 449 | T → R | G domain | | (3) |

| | | | | |
|---|---|---|---|---|
| 459 | G → E | G domain | | (21) |
| 463 | I → IFIF | G domain | | |
| 468 | K → E | G domain | | (4) |
| 470 | D → G | G domain | | (5) |
| 487 | E → K | G domain | In OPA1 and BEHRS. | (3, 14) |
| 503 | T → K | G domain | | (2, 8) |
| 505 | K → N | G domain | | (8) |
| 545 | S → R | Middle | In DOA+ and OPA1; decreased GTPase activity; loss of function in promoting mitochondrial fusion. | (3, 10, 13, 18, 22) |
| 551 | C → Y | Middle | In OPA1 and DOA+. | (3, 23) |
| 551 | Missing | Middle | | (4) |
| 571 | R → H | Middle | | (2) |
| 574 | L → P | Middle | | (5) |
| 586 – 589 | Missing | Middle | | (2) |
| 590 | R → Q | Middle | | (3) |
| 590 | R → W | Middle | | (11) |
| 593 | L → P | Middle | | (3) |
| 593 | Missing | Middle | | (24) |
| 646 | S → L | Middle | | (3) |
| 700-701 | Missing | GMB | | (5) |
| 728 | N → K | GMB | Loss of function in promoting mitochondrial fusion. | (10, 11) |
| 768 | G → D | GMB | | (3) |
| 781 | R → W | GMB | | (3) |
| 785 | Q → R | GMB | Loss of lipid binding and partial loss of function in promoting mitochondrial fusion. | (4, 6, 10) |
| 823 | S → Y | GMB | | (3) |
| 841 | Y → C | GED | | (1) |
| 882 | R → L | GED | | (3) |
| 887 | L → P | GED | | (3) |
| 910 | Missing | GED | | (21) |
| 932 | R → C | GED | | (3, 25) |
| 939 | L → P | GED | Impairs protein folding; loss of function in promoting mitochondrial fusion. | (6, 10) |
| 949 | L → P | GED | | (3, 18) |
| **Dominant optic atrophy plus syndrome (DOA+)** | | | | |
| 357 | A → T | G domain | In DOA+ and OPA1. | (3, 13) |
| 439 | G → V | G domain | In DOA+ and OPA1; decreased GTPase activity; loss of function in | (10, 13, 17) |

| | | | | |
|---|---|---|---|---|
| | | | promoting mitochondrial fusion | |
| 445 | R → H | G domain | In DOA+ and OPA1; decreased GTPase activity; loss of function in promoting mitochondrial fusion | (10, 13, 18-20) |
| 449 | T → P | G domain | | (26) |
| 545 | S → R | Middle | In DOA+ and OPA1; decreased GTPase activity; loss of function in promoting mitochondrial fusion. | (3, 10, 13, 18, 22) |
| 551 | C → Y | Middle | In OPA1 and DOA+. | (3, 23) |
| 582 | Y → C | Middle | | (26) |
| 910 | V → D | GED | Impairs protein folding; loss of function in promoting mitochondrial fusion. | (10, 13) |
| **Behr syndrome (BEHRS)** | | | | |
| 382 | I → M | G domain | In OPA1 and BEHRS | (3, 14, 15) |
| 402 | V → M | G domain | In BEHRSI. | (27) |
| 487 | E → K | G domain | In OPA1 and BEHRS. | (3, 27) |
| **Mitochondrial DNA depletion syndrome 14, cardioencephalomyopathic type (MTDPS14)** | | | | |
| 534 | L → R | Middle | | (28) |
